# FASN-deficiency induces a cytosol-to-mitochondria citrate flux to mitigate detachment-induced oxidative stress

**DOI:** 10.1101/2023.03.14.532533

**Authors:** Wenting Dai, Zhichao Wang, Guan Wang, Qiong A. Wang, Ralph DeBerardinis, Lei Jiang

## Abstract

Fatty acid synthase (FASN) maintains *de novo* lipogenesis (DNL) to support rapid growth in most proliferating cancer cells. Lipogenic acetyl-CoA is primarily produced from carbohydrates but can arise from glutamine-dependent reductive carboxylation under hypoxia. Here we show that reductive carboxylation also occurs in the absence of DNL in cells with defective FASN. In this state, reductive carboxylation was mainly catalyzed by isocitrate dehydrogenase-1 (IDH1) in the cytosol, but IDH1-generated citrate was not used for DNL. Metabolic flux analysis (MFA) revealed that FASN-deficiency induced a net cytosol-to-mitochondria citrate flux through citrate transport protein (CTP). A similar pathway was previously shown to mitigate detachment-induced mitochondrial reactive oxygen species (mtROS) in anchorage-independent tumor spheroids. We further demonstrate that FASN-deficient cells acquire resistance to oxidative stress in a CTP- and IDH1-dependent manner. Together with the reduced FASN activity in tumor spheroids, these data indicate that anchorage-independent malignant cells trade FASN-supported rapid growth for a cytosol-to-mitochondria citrate flux to gain redox capacity against detachment-induced oxidative stress.

## Introduction

Fatty acid metabolism remodeling is essential in the metabolically dynamic tumor microenvironment to support tumorigenesis and cancer progression. Accumulated evidence indicates that proliferating and metastatic cancer cells acquire distinct lipid metabolism to support their growth needs and adapt to environmental challenges (Bergers & Fendt, 2021, Faubert et al., 2020). We previously found that adaptation to anchorage-independence requires a fundamental metabolic change in the citrate metabolism, a lipogenic precursor for *de novo* lipogenesis (DNL), via reductive carboxylation to suppress oxidative stress (Jiang et al., 2016). However, the role of fatty acid synthase (FASN), a critical lipogenic enzyme, in reductive carboxylation during metastatic progression is unclear.

FASN catalyzes an NADPH-dependent reaction to produce fatty acids, which are further utilized for DNL (Röhrig & Schulze, 2016). Proliferating cancer cells often express a high level of FASN to maintain the enhanced membrane synthesis, and FASN inhibition effectively reduces tumor growth in many experimental models (Bandyopadhyay et al., 2006, Bueno et al., 2019, Ferraro et al., 2021). Thus, multiple pharmacologic FASN inhibitors have been developed and applied to preclinical cancer research. However, FASN inhibitors have demonstrated limited clinical success and are not approved for cancer treatment yet (Batchuluun et al., 2022).

FASN utilizes acetyl-CoA to produce fatty acids, and ATP citrate lyase (ACLY) produces lipogenic acetyl-CoA from cytosolic citrate (Droin et al., 2021). Citrate is mainly exported out of mitochondria into the cytosol through citrate transport protein (CTP) in most cells (Arnold et al., 2022, Spinelli & Haigis, 2018, Tan et al., 2020), and carbohydrates are the primary source of lipogenic acetyl-CoA. However, reductive carboxylation serves as an alternative pathway to directly produce citrate from α-ketoglutarate (αKG) derived from glutamine in cancer cells under hypoxia (Metallo et al., 2011). Thus, the primary function of glutamine-dependent reductive carboxylation in cancer cells has been viewed to support DNL (Altman et al., 2016, Metallo et al., 2011, Mullen et al., 2011, Wise et al., 2011).

This study shows that glutamine-dependent reductive carboxylation unexpectedly occurred in the absence of DNL under hypoxia, and FASN-inhibition activates reductive carboxylation under normoxia. This DNL-independent reductive carboxylation was mainly catalyzed by the cytosolic isocitrate dehydrogenase-1 (IDH1), although IDH1-generated cytosolic citrate was not utilized for DNL in FASN-deficient cells. Metabolic flux analysis (MFA) revealed that FASN-deficient cells acquire a net cytosol-to-mitochondria citrate flux through CTP, which is best known to deliver citrate to the cytosol through exchanging mitochondrial citrate with cytosolic malate (Palmieri et al., 2020). FASN-deficient cells depended on this reversed citrate traffic to mitigate detachment-induced oxidative stress. Moreover, while anchorage-independence is a critical characteristic of metastatic cancer cells, FASN was less active in the anchorage-independent tumor spheroids. Taken together, our data suggest that malignant cancer cells reduce FASN-mediated DNL to gain higher redox capacity against external oxidative stress. Thus, this IDH1- and CTP-mediated cytosol-to-mitochondria citrate flux is a metabolic vulnerability in malignant cancer cells, which can potentially serve as therapeutic targets.

## Results

### Reductive carboxylation occurs in the absence of DNL

As illustrated in Figure 1A, previous [U-^13^C]glutamine tracing studies revealed that reductive carboxylation generates citrate to supply lipogenic acetyl-CoA for DNL in hypoxia-cultured cancer cells (Metallo et al., 2011, Wise et al., 2011). It is worth noting that reductive carboxylation also occurs in cancer cells cultured under normoxia, although it remains unclear whether DNL is required for glutamine-dependent reductive carboxylation under normoxia. To test this, we performed [U-^13^C]glutamine tracing in H460 cells incubated with FASN inhibitors C75 and GSK2194069 (Alli et al., 2005, Hardwicke et al., 2014). Both inhibitors reduced intracellular palmitate levels by over 50% (Figure 1B). However, similar to DMSO control, intracellular palmitate remained as ^13^C enriched (containing ^13^Carbon from the [U-^13^C]glutamine tracer) in the C75-treated cells, indicating C75 only partially blocked FASN activity at the dose of 50 µM (Figure 1B). In comparison, 100% of the palmitate was unenriched (m+0, containing 0 ^13^Carbon from [U-^13^C]glutamine tracer) in the GSK2194069-treated cells, showing that GSK2194069 completely blocked FASN activity (Figure 1B). Thus, GSK2194069 was selected to further study the role of FASN in reductive carboxylation.

**Figure 1.**
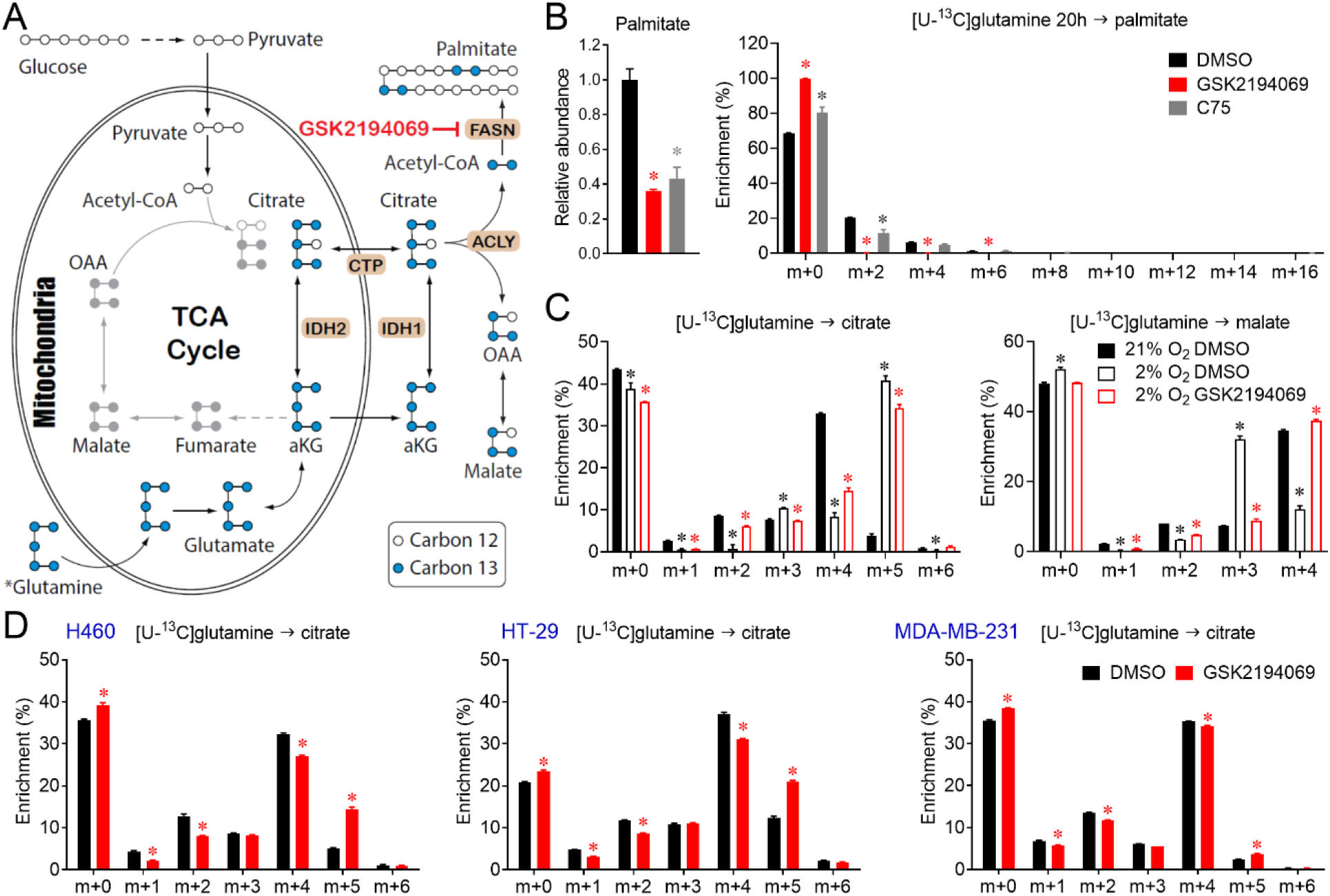
Reductive carboxylation occurs in the absence of *de novo* lipogenesis (DNL). (A) An illustration of the metabolic flow of TCA cycle intermediates and path of citrate to palmitate in cells cultured with [U-^13^C]glutamine under hypoxia. (B) The relative palmitate abundance and mass isotopologue analysis of palmitate enrichment in H460 cells incubated with [U-^13^C]glutamine tracing medium containing DMSO, 50 µM GSK2194069, and 50 µM C75 for 20 hours under normoxia. The mean ± SD error bars are displayed. *P* values are derived from a one-way ANOVA followed by Dunnett’s multiple comparisons test for the three treatments (\**P* < 0.05 compared to the DMSO group). (C) Mass isotopologue analysis of citrate and malate enrichment in H460 cells pretreated with DMSO and 50 µM GSK2194069 for 16 hours and then incubated with the corresponding [U-^13^C]glutamine tracing medium for another 4 hours under normoxia and hypoxia (21% *vs*. 2% O_2_). The mean ± SD error bars are displayed. *P* values are derived from a one-way ANOVA followed by Dunnett’s multiple comparisons test for the three groups (\**P* < 0.05 compared to the 21% O_2__DMSO group). (D) Mass isotopologue analysis of citrate enrichment in multiple cancer cell lines (H460, HT29, and MDA-MB-231) pretreated with DMSO and GSK2194069 for 16 hours and then incubated with [U-^13^C]glutamine tracing medium for another 4 hours. The mean ± SD error bars are displayed. *P* values are derived from a two-tailed Welch’s unequal variances t-test between DMSO- and GSK2194069-treated groups in each cell line (\**P* < 0.05 compared to DMSO treatment). All experiments were repeated 3 times or more, and n = 3 independent samples.

Consistent with the previous studies, compared to normoxia (21% O_2_), hypoxia (2% O_2_) induced reductive labeling in H460 cells as demonstrated by the much higher m+5 labeling of citrate and m+3 labeling of malate (Figure 1C). As illustrated in Figure 1A, m+5 labeling of citrate supplies lipogenic acetyl CoA, m+5 labeling of citrate can be catalyzed cleaved by ACLY into m+3 labeling of malate and m+2 acetyl CoA, which supplies FASN-catalyzed *de novo* fatty acid synthesis. Interestingly, although FASN inhibitor (GSK2194069) blocked the hypoxia-induced m+3 labeling of malate, it only slightly reduced the m+5 labeling of citrate in H460 cells cultured under hypoxia (Figure 1C). These data show that reductive glutamine metabolism persists even when cells do not need this pathway to supply lipogenic acetyl-CoA in the hypoxic condition. To further explore the disconnection between fatty acid synthesis and reductive carboxylation, multiple [U-^13^C]glutamine-cultured cancer cell lines (lung cancer H460, colorectal cancer HT29, and breast cancer MDA-MB-231) were treated with GSK2194069 under normoxia. As in the hypoxia-cultured H460, GSK2194069 reduced m+3 labeling of malate in normoxia-cultured H460 and MDA-MB-231 cells, as well as m+3 labeling of αKG in all three cancer cell lines (Figure S1). Unexpectedly, in all three cell lines, GSK2194069 increased m+5 labeling of citrate from [U-^13^C]glutamine tracer (Figure 1D). These data indicate that reductive carboxylation occurs in the absence of DNL under hypoxia, and FASN inhibition induces reductive carboxylation under normoxia.

### FASN-deficiency enhances reductive carboxylation

To further explore DNL-independent reductive carboxylation, we generated FASN-deficient H460 cells through CRISPR/Cas9 gene editing. Immunoblots showed the complete loss of FASN protein in two pools of FASN-knockout clones (FASN-KO1 and FASN-KO2) (Figure 2A). [U-^13^C]glutamine tracing further confirmed that FASN-KO cells are devoid of *de novo* fatty acid synthesis, although total palmitate level only decreased by 25% (Figure 2B). This suggested that FASN deficiency induced compensatory fatty acids uptake, as the culture medium contained fatty acids from the fetal bovine serum (FBS). FASN deficiency also resulted in the accumulation of NADPH (a substrate of FASN) and the reduction of glutathione (GSH) (Figure 2C and Figure S2A). Additionally, FASN deficiency significantly impaired proliferation in monolayer cultured H460 cells, as well as the spheroid formation under the non-adherent culture (Figure 2D). Together, these data confirm that FASN is essential for the rapid growth of cancer cells, consistent with previous studies (Bueno et al., 2019, Ferraro et al., 2021).

**Figure 2.**
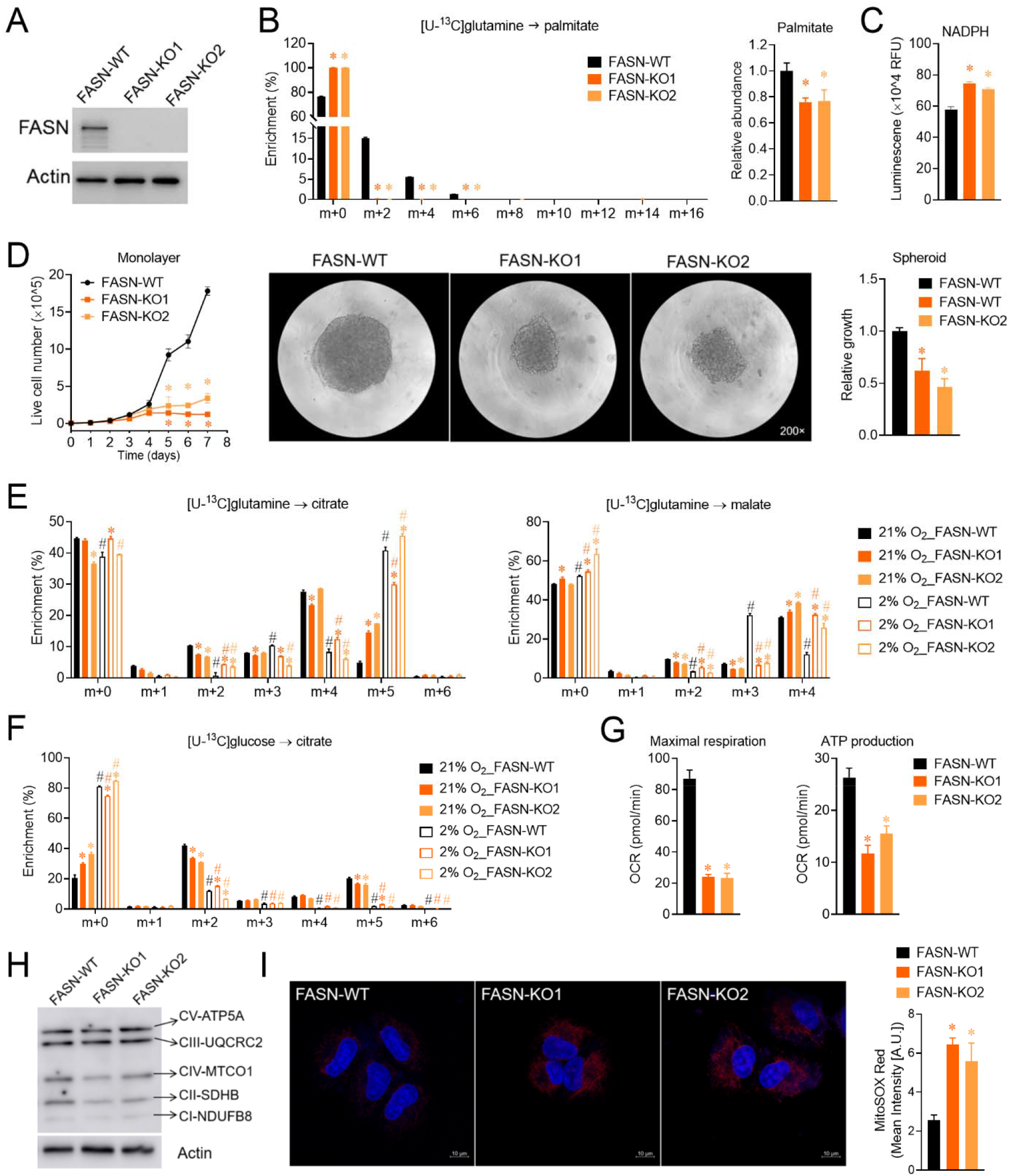
FASN-deficiency enhances reductive carboxylation. (A) Immunoblots of FASN in FASN-wild-type/knockout (FASN-WT/-KO) H460 cells. (B) The relative palmitate abundance and mass isotopologue analysis of palmitate enrichment in FASN-WT and FASN-KO cells incubated with [U-^13^C]glutamine tracing medium for 20 hours. C) The relative abundance of NADPH in FASN-WT and FASN-KO cells. (D) Cell proliferation (n = 3) and spheroid growth (n = 4) of FASN-WT and FASN-KO cells. The spheroid growth of FASN-WT and FASN-KO cells after a 48-hour incubation in non-adherent round bottom 96-well plates (n = 4). The magnification is 200-fold. B-D: The mean ± SD error bars are displayed. *P* values are derived from a one-way ANOVA followed by Dunnett’s multiple comparisons test for the three cell lines (\**P* < 0.05 compared to the FASN-WT group). (E) Mass isotopologue analysis of citrate and malate enrichment in FASN-WT and FASN-KO cells incubated with medium containing [U-^13^C]glutamine tracing medium for 4 hours under normoxia and hypoxia (21% *vs*. 2% O_2_). (F) Mass isotopologue analysis of citrate in FASN-WT and FASN-KO H460 cells incubated with [U-^13^C]glucose tracing medium for 4 hours under normoxia and hypoxia (21% *vs*. 2% O2). E and F (n = 3): the mean ± SD error bars are displayed. *\*P* values are derived from a one-way ANOVA followed by Dunnett’s multiple comparisons test for the three cell lines under normoxia or hypoxia (\**P* < 0.05 compared to 21% O_2__FASN-WT group or 2% O_2__FASN-WT group). ^#^*P* values are derived from a two-tailed Welch’s unequal variances t-test between two groups in each cell line (^#^*P* < 0.05 compared 21% O_2_ group to 2% O_2_ group). (G) The maximal respiration and ATP production in FASN-WT and FASN-KO H460 cells (n = 6– 7). The mean ± SD error bars are displayed. P values are derived from a one-way ANOVA followed by Dunnett’s multiple comparisons test for the three cell lines (*P < 0.05 compared to the FASN-WT group). (H) Immunoblots of electron respiration chain (ETC) complexes subunits in FASN-WT and FASN-KO cells. ETC I, NDUFB8, NADH dehydrogenase [ubiquinone] 1 beta subcomplex subunit 8; ETC II, SDHB, succinate dehydrogenase [ubiquinone] iron-sulfur subunit b; ETC III, UQCRC2, cytochrome b-c1 complex subunit 2; ETC IV, MTCO1, cytochrome c oxidase subunit I; ETC V, ATP5A, ATP synthase lipid-binding protein. (I) Mitochondrial ROS (mtROS) levels were measured by mitoSOX red staining in FASN-WT and FASN-KO cells (n = 4). The scale bar represents 10 μm. The mean ± SD error bars are displayed. *P* values are derived from a one-way ANOVA followed by Dunnett’s multiple comparisons test for the three treatments (\**P* < 0.05 compared to the FASN-WT group). All experiments were repeated 3 times or more.

Under normoxia, FASN-KO cells also exhibited higher m+5 labeling of citrate than FASN-WT H460 cells (Figure 2E), consistent with the elevated reductive carboxylation in the GSK2194069-treated cells. Under hypoxia, FASN-WT and FASN-KO cells had a similarly high level of reductive carboxylation, which produced about 40% of the total intracellular citrate. Additionally, FASN-deficiency reduced m+3 labeling of malate, fumarate, and aspartate under both normoxia and hypoxia, while there was not much change in m+5 labeling of αKG and glutamate from [U-^13^C]glutamine in these cells (Figure 2E and Figure S2B). These data suggest that glutamine-dependent reductive carboxylation occurs in FASN-deficient cells regardless of oxygen status, and FASN-deficiency stimulates reductive carboxylation under normoxia.

Biochemically, a higher αKG/citrate (substrate/product) ratio is the best-known driver of the reductive carboxylation of αKG into citrate (Fendt et al., 2013). FASN-KO and FASN-WT had a similar intracellular level of αKG and citrate (Figure S2C), indicating that a higher αKG/citrate is not the mechanism of the elevated glutamine-dependent reductive carboxylation observed in FASN-deficient cancer cells. To further explore how FASN-deficiency induces reductive carboxylation, we assessed mitochondrial oxidative metabolism in FASN-WT and FASN-KO H460 cells under different oxygen levels, as reductive carboxylation is also known to be induced by electron transport chain (ETC)-deficiency and reduced mitochondrial oxidative phosphorylation (Metallo et al., 2011, Mullen et al., 2011, Wise et al., 2011). Under both normoxia and hypoxia, FASN-WT and FASN-KO cells had similar pyruvate and lactate enrichment from [U-^13^C]glucose tracer (Figure S2D). In contrast, only normoxia-cultured FASN-KO cells had lower m+2 labeling of citrate (Figure 2F), while the enrichment of other TCA cycle intermediates (malate, fumarate, and αKG) was similar between FASN-WT and FASN-KO cells (Figure S2E). Consistent with the repressed glucose oxidation, the maximal respiration and ATP production were lower in FASN-KO cells, although the basal respiration was similar between FASN-WT and FASN-KO cells (Figure 2G and Figure S2F). Furthermore, FASN-KO cells expressed lower level of ETC complex II (SDHB, succinate dehydrogenase [ubiquinone] iron-sulfur subunit) and IV (MTCO1, cytochrome c oxidase subunit I), and FASN-KO cells had a higher level of mitochondrial reactive oxygen species (mtROS) (Figure 2H and 2I). Together, these data show that FASN-deficiency represses mitochondrial activity. This is in line with a recent study, when electron acceptors are limited in proliferating cancer cells under hypoxia, these cancer cells rely more extracellular lipids to preserve mitochondrial activity (Li et al., 2022).

To further confirm the roles of FASN in these metabolic alterations, we genetically rescued FASN expression in FASN deficient cell through lentivirus-mediated FASN overexpression (Figure 3A). As expected, FASN overexpression rescued the impaired growth in monolayer cultured cells, as well as the spheroid size under the non-adherent condition (Figure 3B). FASN overexpression also reduced mtROS in FASN deficient cell (Figure 3C). Additionally, FASN overexpression also blocked the induced reductive carboxylation in the FASN deficient cell (Figure 3D). Together, these data show that FASN deficiency enhances glutamine dependent reductive carboxylation.

**Figure 3.**
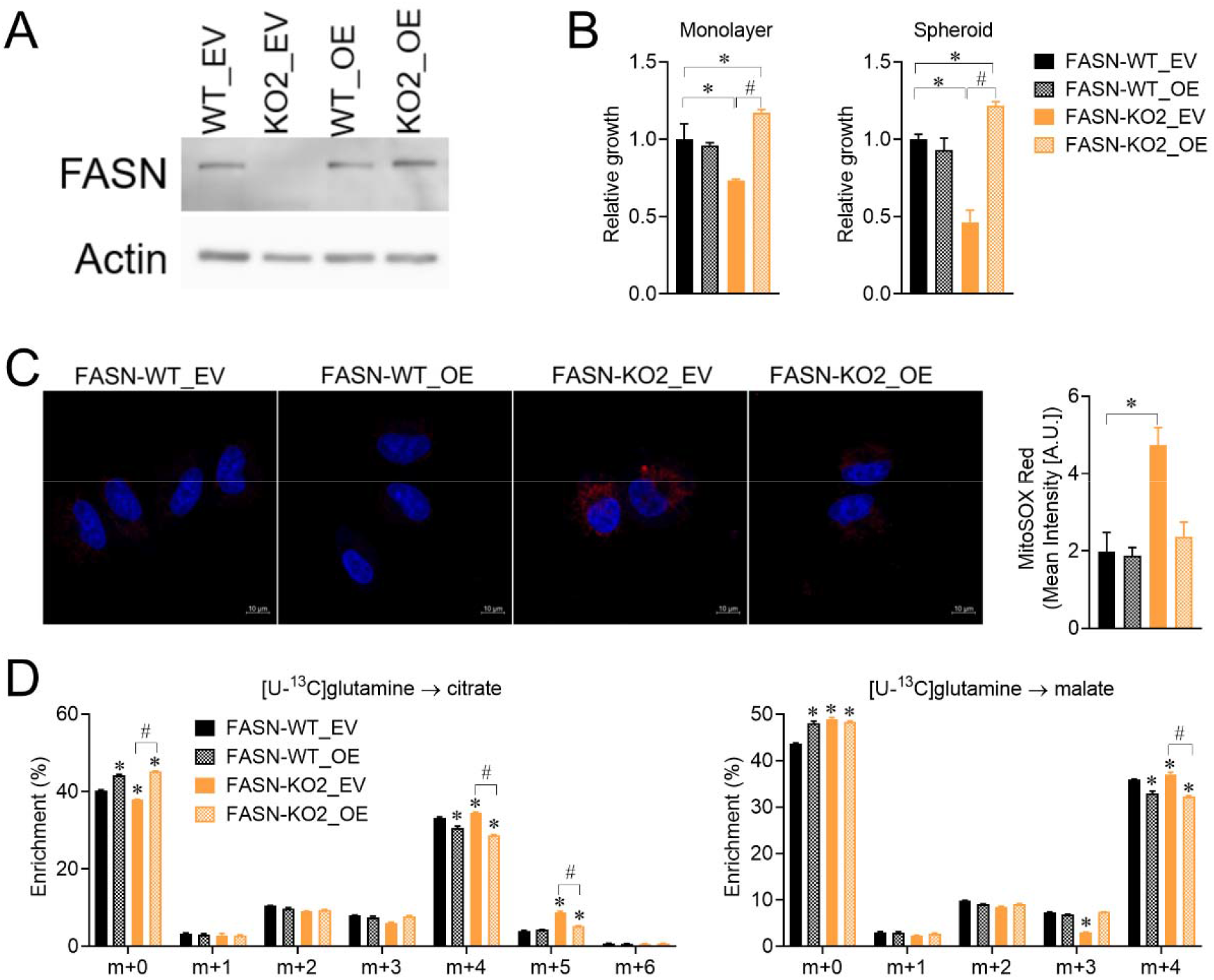
FASN overexpression reduces reductive carboxylation in FASN-KO cells (Supporting data for Figure 2). FASN-WT and FASN-KO2 cells were transfected with FASN overexpression (OE) or empty vector (EV) plasmid. (A) Immunoblots of FASN and β-actin. (B) Relative growth of monolayer and spheroid after a 24-hour incubation in attached flat-bottom and non-adherent round-bottom 96-well plates, respectively (n= 4). (C) Mitochondrial ROS levels measured by mitoSOX Red staining (n = 4). The scale bar represents 10 μm. (D) Mass isotopologue analysis of citrate and malate enrichment in cells incubated with medium containing [U-^13^C]glutamine tracing medium for 4 hours (n = 3). B-D: The mean ± SD error bars are displayed. P values are derived from a two-way ANOVA followed by Dunnett’s multiple comparisons test for the four groups (\**P* < 0.05 compared to the FASN-WT_EV group, ^#^*P* < 0.05 compared to the FASN-KO2_EV group). All experiments were repeated three times or more.

### FASN-deficiency drives a net cytosol-to-mitochondria citrate flux

We previously reported that reductive carboxylation is induced in anchorage-independent H460 spheroids, which also have a higher level of mtROS (Jiang et al., 2016). Thus, we tested whether the induction of reductive carboxylation was dependent on the high mtROS induction in the FASN-deficient cells. First, we used DMNQ, a membrane-permeable redox-cycling quinone, to specifically induce the mtROS levels in FASN-WT and FASN-KO cells (Figure S3A). [U-^13^C]glutamine tracing showed that DMNQ increased the relative m+5 enrichment of citrate in both FASN-WT and FASN-KO cells (Figure S3B). It is worth noting that, since DMNQ significantly reduced the total level of intracellular citrate, DMNQ did not alter the absolute level of m+5 citrate in either FASN-WT or FASN-KO cells when the relative abundance of each citrate isotopologue was calculated (Figure S3C). To further explore the causal relation between mtROS and reductive carboxylation, we also used mitoTEMPO to specifically reduce mtROS levels in FASN-KO cells (Figure S3A). [U-^13^C]glutamine tracing showed that mitoTEMPO-treated FASN-KO cells maintained their reductive carboxylation (Figure S3D). Collectively, these results suggest that mtROS and reductive carboxylation are concurrently induced in the FASN-deficient cells.

A pair of IDH isoforms, IDH1 and IDH2, catalyze reductive carboxylation in the cytosol and mitochondria, respectively (Jiang et al., 2016, Metallo et al., 2011). To test which IDH isoform mediates the reductive carboxylation in FASN-deficient cells, we performed [U-^13^C]glutamine tracing in FASN-WT and FASN-KO cells in the presence of IDH1 inhibitor (IDH1i, GSK321), IDH2 inhibitor (IDH2i, AGI-6780), or both IDH1/2 inhibitors. IDH1 inhibition alone blocked most (80%) m+5 labeling of citrate in FASN-WT and FASN-KO cells, while IDH2 inhibition alone slightly (20%) reduced m+5 labeling of citrate in FASN-KO cells (Figure 4A). Moreover, the combined inhibition of IDH1 and IDH2 resulted in a complete loss (up to 95%) of m+5 labeling of citrate in both FASN-WT and FASN-KO cells. In contrast to citrate enrichment, although IDH1 inhibitor partially reduced m+3 labeling of malate in FASN-WT cells, neither IDH1/2 inhibitor alone nor in combination altered m+3 labeling of malate in FASN-KO cells (Figure 4B); this is expected, as reductive carboxylation only contributes to lipogenic acetyl-CoA in FASN-WT cells but not in the FASN-KO cells. These data suggest that reductive carboxylation is mainly catalyzed by the cytosolic IDH1 in FASN-deficient cells, although it is somewhat surprising that IDH1-generated cytosolic citrate is not utilized for DNL.

**Figure 4.**
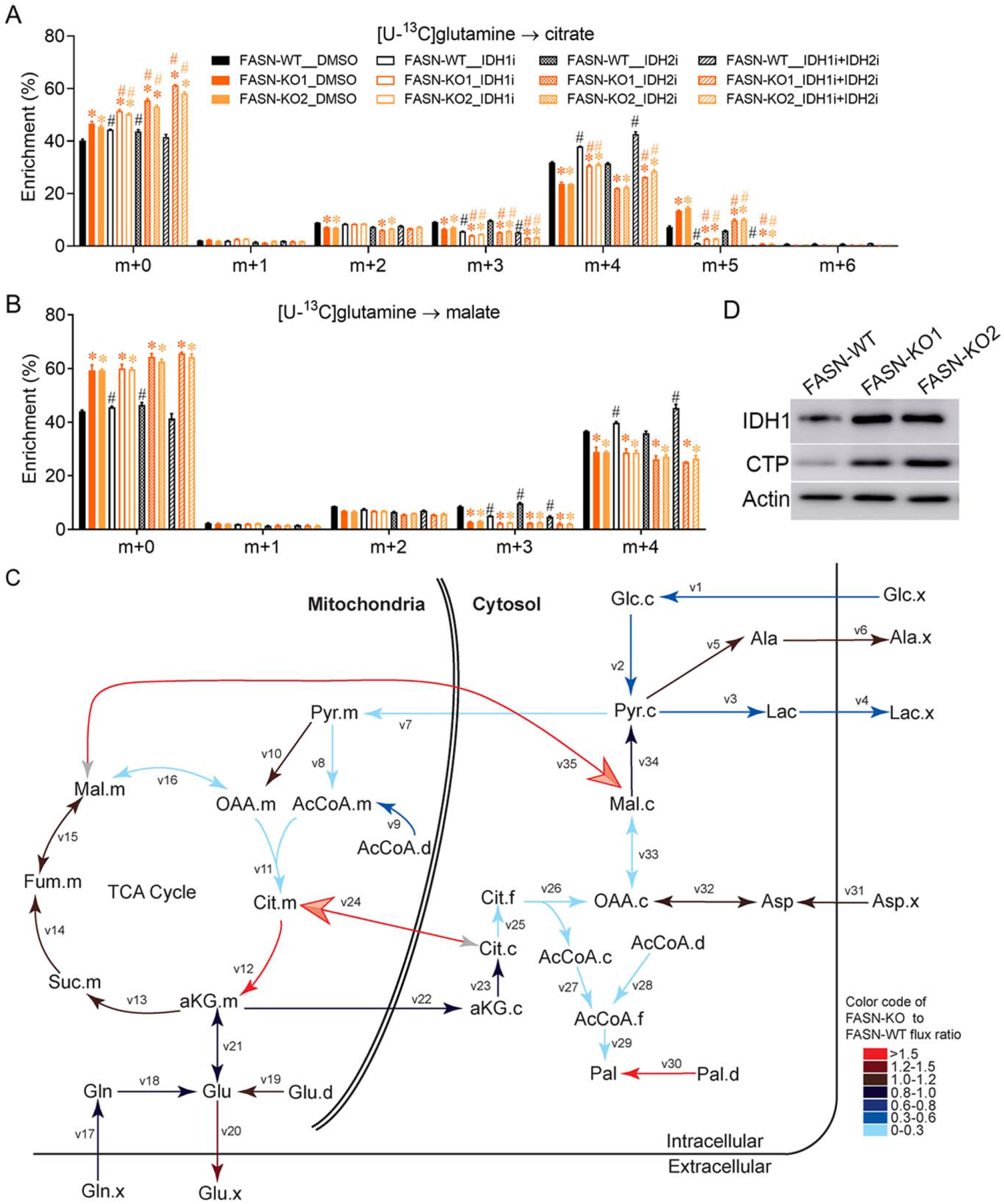
FASN deficiency drives a net cytosol-to-mitochondria citrate flux. (A) and (B) Mass isotopologue analysis of citrate and malate enrichment in FASN-WT and FASN-KO cells incubated with [U-^13^C]glutamine tracing medium containing 5 µM IDH1 inhibitor (IDH1i, GSK321), 5 µM IDH2 inhibitor (IDH2i, AGI-6780), and IDH1i plus IDH2i for 4 hours. The mean ± SD error bars are displayed. *\*P* values are derived from a one-way ANOVA followed by Dunnett’s multiple comparisons test for the three cell lines under the same treatment (\**P* < 0.05 compared to FASN-WT, ^#^*P* < 0.05 compared to FASN-WT/-KO_DMSO group). A-B: The experiments were repeated 3 times or more, and n = 3 independent samples. (C) The modeling of metabolic fluxes in FASN-deficient cells. The comparative modeling of metabolic fluxes in FASN-KO2 H460 cells to FASN-WT controls. See **Table S1** for definitions of quantitative flux values. The color scale reflects the ratio of each flux in FASN-KO cells relative to FASN-WT controls. (D) Immunoblots of IDH1, CTP, and β-actin in FASN-WT and FASN-KO cells. IDH1, isocitrate dehydrogenase 1; CTP, citrate transport protein.

To better understand the disconnection between the absence of DNL and the induction of reductive carboxylation, we next performed MFA to systemically interpret the metabolic reprogramming in the FASN-deficient cells. The reaction network describing the stoichiometry and carbon transitions of central carbon metabolism was modified from our previously MFA modeling of anchorage-independent tumor spheroids (Table S1) (Jiang et al., 2016). Steady-state fluxes were calculated by integrating extracellular flux rates and ^13^C distributions in several metabolites from both [U-^13^C]glucose and [U-^13^C]glutamine tracers. Modeling confirmed that the ratio of reductive to oxidative citrate production was much higher in FASN-KO cells, compared to FASN-WT cells (Table S1). Moreover, modeling showed that IDH1-generated citrate entered mitochondria in FASN-KO cells (Figure 4C). Consistent with the MFA modeling, immunoblots showed that FASN-deficient H460 cells expressed a higher level of IDH1 and CTP (Figure 4D). It is worth noting that, although our previously published modeling indicates bidirectional citrate fluxes across the mitochondrial membrane, the net citrate flux in tumor spheroid remains in the usual mitochondria-to-cytosol direction (Jiang et al., 2016). In contrast, the current modeling indicated that FASN-KO cells had a net cytosol-to-mitochondria citrate flux. Together, these findings from the MFA model indicate that reducing FASN activity results in an unusual cytosol-to-mitochondria citrate flux.

### Cytosol-to-mitochondria citrate flux protects FASN-deficient cells against detachment-induced oxidative stress

Quantitative MFA indicates that FASN-deficient cells and anchorage-independent tumor spheroids share similar metabolic reprogramming, an induction of cytosolic reductive carboxylation followed by an unusual cytosol-to-mitochondria citrate flux. Interestingly, by re-analyzing the palmitate enrichment data of [U-^13^C]glutamine and [U-^13^C]glucose cultured H460 tumor spheroids (Jiang et al., 2016), we found that the palmitate synthesis rate was lower in anchorage-independent tumor spheroids compared to monolayer culture (Figure S4A). We also analyzed the data from ctcRbase, which is an online database to collect expression data of circulating tumor cells (CTC) and primary tumors from cancer patients (Gene expression database of CTC). RNA-seq data analyses showed that CTC had lower FASN expression levels, compared to their primary tumor cells in breast cancer (Yu et al., 2014) and prostate cancer (Miyamoto et al., 2015) (Figure S4B). These data suggest that FASN is less active in the cultured anchorage-independent spheroids and in the cancer patients CTC, which are in line with the compromised proliferation of matrix-detached cells (Grassian et al., 2011).

While reductive carboxylation and cytosol-to-mitochondria citrate flux mitigate oxidative stress in anchorage-independent tumor spheroids (Jiang et al., 2016), adherent FASN-deficient cells had a higher basal level of mtROS than FASN-WT cells (Figure 2I). We next tested how detached FASN-deficient and FASN-expressing cells respond to external oxidative stress. The mitoSOX staining showed that DMNQ increased mtROS in detached FASN-WT cells, to a much higher extent, than that in FASN-KO cells (Figure 5A). When detached cells were treated with DMNQ, the relative growth was reduced to 30% of the DMSO control in FASN-WT cells versus 60∼70% of the DMSO control in FASN-KO cells, suggesting detached FASN-WT cells were more sensitive to external oxidative stress (Figure 5B). Moreover, when the cytosol-to-mitochondria citrate flux was blocked by CTP inhibitor (CTPi), IDH1 inhibitor (IDH1i), or IDH2 inhibitor (IDH2i), DMNQ co-treated FASN-KO2 cells showed higher mtROS than DMNQ-treated FASN-KO2 cells (Figure 5C). In line with the changes in mtROS, the relative growth of DMNQ co-treated FASN-KO2 cells under detachment were further depressed compared to DMNQ-treated FASN-KO2 cells (Figure 5D). Together, these data show that FASN-deficient cells rely on cytosol-to-mitochondria citrate flux to mitigate detachment- and DMNQ-induced oxidative stress.

**Figure 5.**
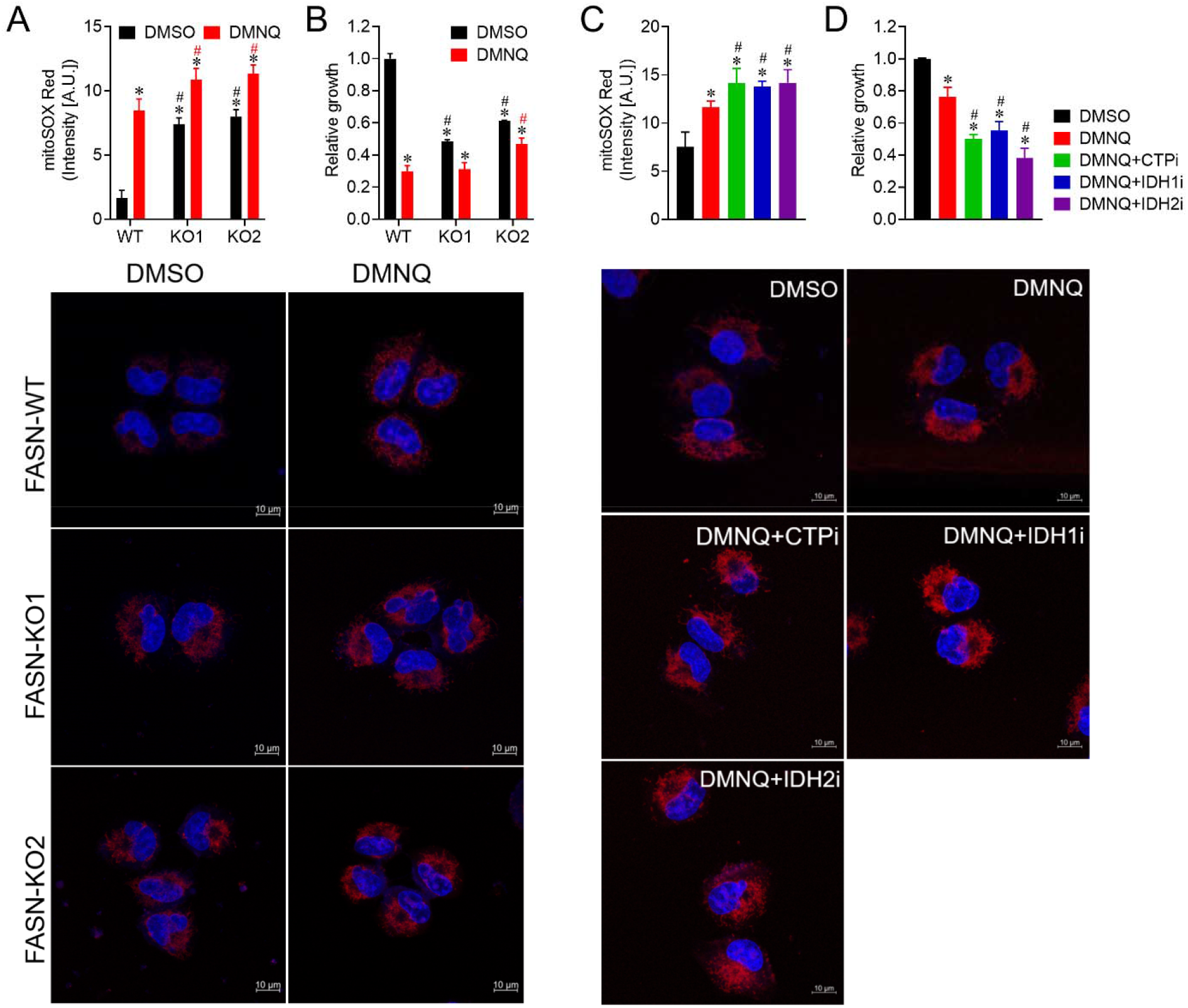
Cytosol-to-mitochondria citrate flux protects FASN-deficient cells against detachment-induced oxidative stress. (A) Mitochondrial ROS levels measured by mitoSOX Red staining and their quantification in detached FASN-WT and FASN-KO cells exposure to DMSO and 10 µM DMNQ for 24 hours (n = 4). (B) Relative growth of detached FASN-WT and FASN-KO cells exposure to DMSO and 10 µM DMNQ for 24 hours (n = 4). (C) Mitochondrial ROS levels measured by mitoSOX Red staining and their quantification in detached FASN-KO2 cells exposure to DMSO, 10 µM DMNQ, and 10 µM DMNQ plus 5 µM IDH1i/5 µMIDH2i/200 µM CTPi for 24 hours (n = 4). (D) Relative growth of detached FASN-KO2 cells exposure to DMSO, 10 µM DMNQ, and 10 µM DMNQ plus 5 µM IDH1i/5 µMIDH2i/200 µM CTPi for 24 hours (n = 4). A and C: the scale bar represents 10 μm. A-D: The mean ± SD error bars are displayed. *\*P* values are derived from a one-way ANOVA followed by Dunnett’s multiple comparisons test for the three cell lines under the same treatment (A and B: \**P* < 0.05 compared to FASN-WT_DMSO, ^#^*P* < 0.05 compared to FASN-WT/-KO_DMSO group in each line; C and D: \**P* < 0.05 compared to DMSO group, ^#^*P* < 0.05 compared to DMNQ group). All experiments were repeated 3 times or more.

## Discussion

This study shows that reductive carboxylation occurs in the absence of DNL under hypoxia, and FASN inhibition induces reductive carboxylation under normoxia. FASN inhibition has been viewed as an effective approach to limit tumor growth since most proliferating cancer cells rely on active DNL to generate new membranes for daughter cells (Alli et al., 2005, Bueno et al., 2019, Ferraro et al., 2021). However, to date, none of the FASN inhibitors has been approved for cancer treatment (Batchuluun et al., 2022). Our stable isotope tracing assays show that GSK2194069, as a FASN inhibitor, completely blocks *de novo* fatty acids synthesis in multiple cancer cell lines. Additionally, orlistat, a lipase and FDA-approved drug to treat obesity, has also been reported to inhibit FASN (Knowles et al., 2004, Kridel et al., 2004, Zhi et al., 1999). We confirmed that orlistat completely inhibited *de novo* fatty acid synthesis (Figure S5A), and orlistat treatment impaired cell proliferation in H460 cells (Figure S5B). Like GSK2194069, orlistat also induced reductive carboxylation in H460 cells (Figure S5C). Altogether, FASN inhibitors (GSK2194069 and orlistat) promote DNL-independent reductive carboxylation in cancer cells under normoxia.

Consistent with the cancer cells treated with FASN-inhibitors, FASN-deficient H460 lung cancer cells also have a higher level of reductive carboxylation than the FASN-expressing cells. [U-^13^C]glutamine tracing showed that FASN-deficiency increased m+5 labeling of citrate but reduced m+3 labeling of malate. In contrast, hypoxia induces both m+5 labeling of citrate and m+3 labeling of malate synthesis (Metallo et al., 2011, Wise et al., 2011), as illustrated in Figure 1A. The difference in m+3 labeling of malate shows that glutamine-derived m+5 labeling of citrate is not cleaved by ACLY for lipogenic acetyl-CoA production in FASN-deficient cells. Unexpectedly, reductive carboxylation in FASN-deficient cells is mainly catalyzed by IDH1 to generate cytosolic citrate, although it is not utilized for DNL. Moreover, MFA modeling indicates that IDH1-generated cytosolic citrate enters mitochondria through CTP in FASN-deficient cells; this CTP-mediated cytosol-to-mitochondria citrate flux also exists in the anchorage-independent tumor spheroids (Jiang et al., 2016), which may be associated with the CTP-mediated role in sustaining redox homeostasis in cancer cells (Hlouschek et al., 2018). Indeed, anchorage-independent spheroids have less active FASN than monolayer-cultured cells. Together, our studies indicate that anchorage-independent growth represses DNL to induce cytosolic reductive carboxylation followed by an unusual cytosol-to-mitochondria citrate flux.

Anchorage-independence is a property of metastatic cancer cells. Compared to the primary tumors, metastasis (metastatic cancer) is the most common cause of death in cancer patients. While most primary tumors have enhanced anaerobic glycolysis (Warburg effect) (Hanahan, 2022), the more energetically efficient oxidative phosphorylation has been reported in metastatic cancer cells to support their elevated energy need (Fischer et al., 2019). Accumulated evidence indicates that proliferating and metastatic cancer cells prefer distinct energy metabolism, as well as distinct lipid metabolism (Bergers & Fendt, 2021, Faubert et al., 2020). For example, although FASN activation is commonly viewed as a metabolic feature of proliferating cancer cells in primary tumors, both FASN activation and FASN inhibition have been associated with metastatic cancer cells (Jiang et al., 2015a, Jiang et al., 2014, Jiang et al., 2015b, Seguin et al., 2012, Zaytseva et al., 2012).

We previously reported that *de novo* fatty acid synthesis is reduced in the A549 lung cancer cells undergoing epithelial-mesenchymal transition (EMT), and FASN-knockdown increases tumor metastasis *in vivo* (Jiang et al., 2015b). We proposed that FASN-knockdown increases acetyl-CoA supply for mitochondrial oxidative phosphorylation, which increases cellular ATP content and oxygen consumption rate (Jiang et al., 2015a). In the current study, we show that FASN is less active in anchorage-independent tumor spheroids, consistent with MFA in our previous study (Jiang et al., 2016). In contrast, other studies shows that FASN-overexpression increases the migration *in vitro* and experimental peritoneal dissemination *in vivo* in the ovarian cancer (Jiang et al., 2014), and FASN inhibition impairs the metastasis of colorectal cancer and melanomas (Pavlova et al., 2022, Sies et al., 2022). More recently, another isotope tracing study shows that fatty acid synthesis is elevated in breast tumors growing in the brain, where lipid availability is relatively lower than the breast (Faubert et al., 2020). It is worth noting that the gene expression levels of FASN were lower in CTC from breast cancer (Yu et al., 2014) and prostate cancer (Miyamoto et al., 2015) patients compared to their primary cancer cells, which is in line with our finding of FASN-deficiency potentially promoting cancer metastasis. Together, the role of FASN in metastasis is context-dependent, and additional studies are required to determine how to therapeutically target FASN to treat cancer patients.

Moreover, proliferating and metastatic cancer cells also have different redox metabolism, related to their distinct energy metabolism (Pavlova et al., 2022, Sies et al., 2022, Tasdogan et al., 2021). While proliferating cancer cells receive growth and survival stimuli through their attachment to extracellular matrix (ECM), metastatic cancer cells can survive without ECM-induced signals (Valastyan & Weinberg, 2011). Detachment from ECM is associated with enhanced ROS production (Schafer et al., 2009). FASN-deficiency increases intracellular NADPH levels in H460 cells, as the FASN reaction consumes a large amount of cytosolic NADPH, which is also a key co-factor for IDH1-mediated reductive carboxylation (Yoo et al., 2008). Thus, accumulated NADPH in FASN-deficient H460 cells can drive the reductive direction of cytosolic IDH1. Similar to the metabolic reprogramming in H460 spheroids (Jiang et al., 2016), MFA modeling indicates that IDH1-generated citrate enters mitochondria and releases NADPH through IDH2-mediated oxidation, which connects the redox metabolism between cytosol and mitochondria in the FASN-deficient cells. In comparison, FASN-knockdown mouse embryonic fibroblasts (MEFs) expressing PyMT (polyomavirus middle T antigen) unexpectedly have inactive reductive carboxylation despite the presence of NADPH accumulation in these cells (Bueno et al., 2019). The opposite findings in FASN-KO MEFs and H460 cells indicate that reductive carboxylation may be a metabolic feature of metastatic cancer cells, which may mitigate detachment-induced oxidative stress by limiting DNL to reprogram cytosolic and mitochondrial redox metabolism (Figure 6). Together, our studies collectively suggest that targeting cytosol-to-mitochondria citrate flux in the three points can be an effective therapy for treating cancer patients with metastasis.

**Figure 6.**
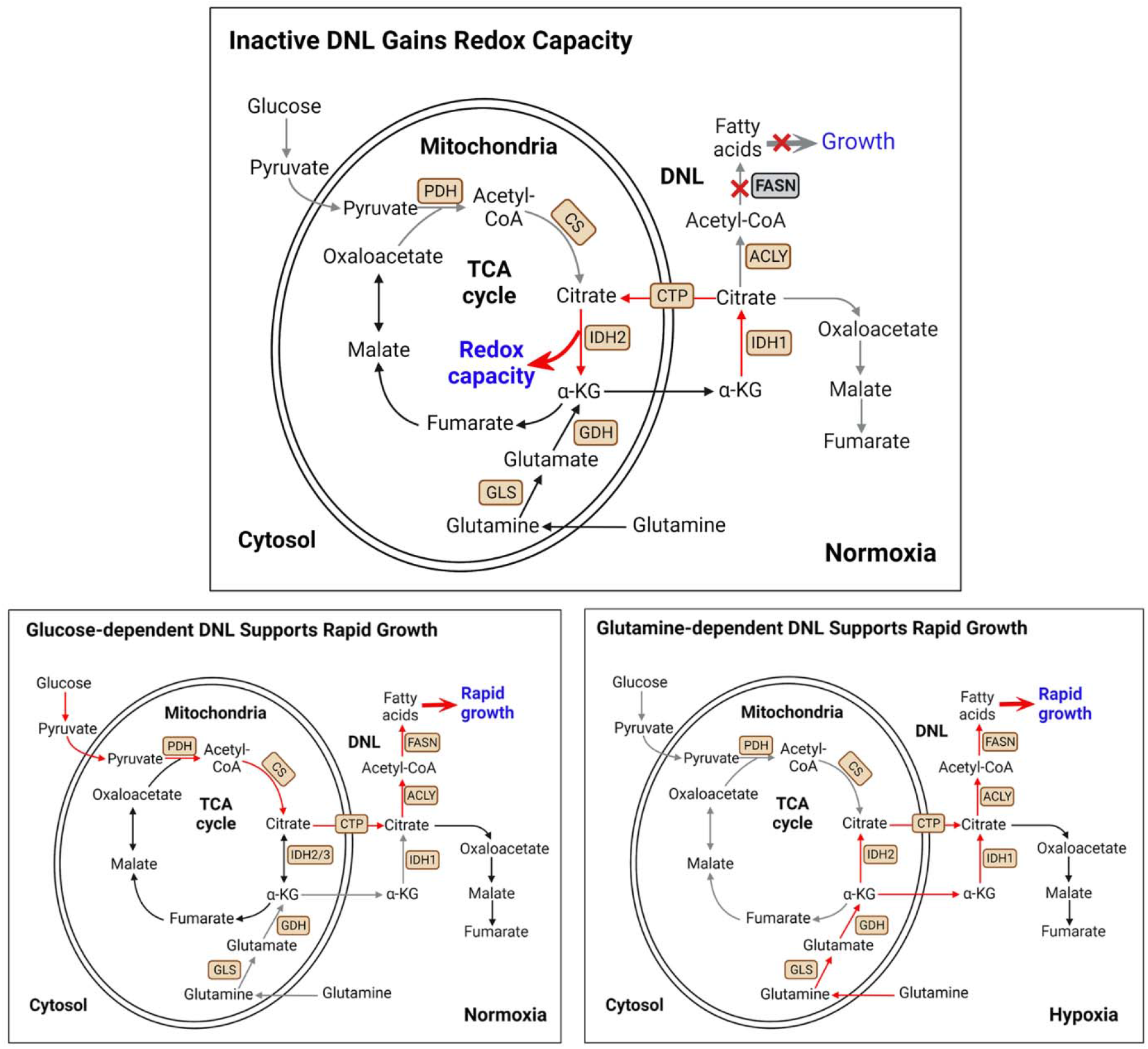
Model illustrates that inactivated DNL leads to the gain of redox capacity. This function goes through the cytosolic IDH1-dependent reductive carboxylation followed by a net cytosol-to-mitochondria citrate flux to mitigate oxidative stress.

## Materials and Methods

### Generation of FASN-knockout (KO) H460 cells

FASN-deficient H460 cells were generated using the CRISPR/Cas9 system (Ran et al., 2013). Wild-type clones were selected from both the control vector and targeting vector transfections. To control the variations among individual clones from single-cell cloning, 4 to 5 clones were pooled together for further experiments (Jiang et al., 2016).

### Generation of genetical FASN-rescued FASN-KO H460 cells

One 10-cm dish of HEK293T cells at 60% confluency were transfected with 6µg psPAX2, 1.5 µg pMD2.G and 7.5 µg pLJM1-Flag-FASN plasmid (FASN-overexpression, FASN-OE) or pLJM1-Empty vector control plasmid (EV) using 15 µL Lipofectamine 3000 diluted in 1mL of OPTI-MEM. Medium containing lentiviral particles was harvested 48 and 72 hours after transfection. Supernatant was filtered using a 0.45µm PVDF filter, divided into aliquots and frozen at -80 °C. The same day, FASN-wildtype (WT)/-KO H460 cells in 6-well plate were transfected by adding 500 µL lentivirus carrying EV and FASN-OE plasmids per well. Addition of polybrene (final concentration of 10 µg/mL) to each well. 24 hours post-transduction, the media of each well were changed with complete culture media. 48 hours post-media change, cells were selected using 5 µg/mL puromycin for one week. The single clone was obtained by serial dilution.

### Cell culture conditions

Lung cancer cell lines H460, H460-FASN-WT/KO, H460-FASN-WT_EV/OE, and H460-FASN-KO2_EV/OE were expanded in RPMI-1640 medium supplemented with 5% fetal bovine serum (FBS; HyClone, CA, USA), 1% penicillin/streptomycin, and 4mM L-glutamine. Colon cancer cell line HT29 and breast cancer cell line were expanded in DMEM medium supplemented with 10% fetal bovine serum (FBS; HyClone, CA, USA), 1% penicillin/streptomycin, and 4mM L-glutamine.

### Stable isotope tracing, sample collection, and analysis

Stable isotope tracing experiments to determine isotope distributions in soluble metabolites were performed as described (Dai et al., 2020). Glucose and glutamine labeled with ^13^C were both purchased from Sigma (MO, USA). Studies of H460 cells (H460, H460-FASN-WT, H460-FASN-KO1/2, H460-FASN-WT_EV/OE, H460-FASN-KO2_EV/OE) were performed in RPMI medium containing 5% dialyzed FBS. HT29 and MDA-MB-231 ^13^C-tracing studies were performed in DMEM medium containing 10% dialyzed FBS. DMEM lacking glucose and glutamine was prepared from powder (Sigma), then supplemented with 10 mM unlabeled glucose and 3 mM [U-^13^C]glutamine. RPMI lacking glucose was prepared from powder (Sigma), and RPMI lacking glutamine was purchased from Sigma, then supplemented with either 10 mM D-[U-^13^C]glucose or 3 mM L-[U-^13^C]glutamine. To explore the effects of FASN inhibitors on palmitate synthesis, we directly incubated cancer cells (H460, HT29, and MDA-MB-231) with the [U-^13^C]glutamine tracing medium containing DMSO or FASN inhibitors, 20/50 μM GSK2194069 (Alli et al., 2005), 50 μM Orlistat, and 50 μM C75 (Alli et al., 2005, Menendez & Lupu, 2007) for 20 hours. To test the effects of FASN inhibitors on cellular metabolism, multiple cancer cell lines (H460, HT29, and MDA-MB-231) were pre-treated with DMSO, 20/50 μM GSK2194069 (or 50 μM Orlistat in H460 cells) for 16 hours followed by another 4 hours of ^13^C-labeled glutamine tracing. To examine the effects of FASN deficiency on palmitate synthesis, FASN-WT and FASN-KO H460 cells were incubated with ^13^C-labeled glutamine medium for 20 hours. To determine the effects of FASN inhibitor on cellular metabolism under hypoxia (2% O_2_) *vs.* normoxia (21% O_2_), H460 cells were grown in 60 mm dishes until 80% confluent, pre-treated with DMSO and 50 μM GSK2194069 for 16 hours, and then cultured with ^13^C-labeled glutamine RPMI medium containing the corresponding treatments for another 4 hours. To examine the effects of FASN deficiency on cellular metabolism in H460 cells under hypoxia *vs.* normoxia, H460-FASN-WT and H460-FASN-KO cells were incubated with ^13^C-labeled glucose and ^13^C-labeled glutamine medium for 4 hours. The hypoxic culture was conducted by feeding a custom mixture of 2% O_2_, 5% CO_2,_ and 94% N_2_ *vs*. a standard incubator controlled at 5% CO_2_. To examine the effects of FASN overexpression on cellular metabolism, H460-FASN-WT_EV/OE and H460-FASN-KO2_EV/OE cells were incubated with ^13^C-labeled glutamine medium for 4 hours.

To investigate whether IDH1 and CTP are involved in FASN deficiency-associated reductive metabolism, H460-FASN-WT and H460-FASN-KO cells were incubated with ^13^C-labeled glutamine medium containing DMSO, 5 µM IDH1 inhibitor (IDH1i, GSK321), or 200 µM CTP inhibitor (CTPi) for 4 hours. To further examine the involvement of IDH1 and IDH2 in FASN-deficiency-associated reductive metabolism, H460-FASN-WT and H460-FASN-KO cells were incubated with ^13^C-labeled glutamine medium containing DMSO, 5 µM IDH1 inhibitor (IDH1i, GSK321), 5 µM IDH2 inhibitor (IDH2i, AGI-6780), and the combination of 5 µM IDH1i and IDH2i for 4 hours. To further examine the effects of mitochondrial ROS level (mtROS) on reductive metabolism, we performed the ^13^C-labeled glutamine tracing containing mtROS inducer (10 µM DMNQ) and mtROS reducer (100 µM mitoTEMPO) for 4 hours in FASN-WT and FASN-KO cells.

The above cells were collected using 50% methanol in water to measure cellular metabolites, as previously described (Wang et al., 2020). Macromolecules and debris were removed by centrifugation (12,000×rpm) for 15 min at 4°C. Subsequently, the supernatants with aqueous metabolites were evaporated, derivatized for 2 hours at 42°C in 50μL of methoxyamine hydrochloride (Sigma) and 100μL N-tert-Butyldimethylsilyl-N-methyltri fluoroacetamide (Sigma, MO, USA) for 90 min at 72°C. Lipids were extracted completely as previously described (Yang et al., 2014). All the metabolites were analyzed using an Agilent 7890B gas chromatograph (Agilent, CA, USA) networked to an Agilent 5977B mass selective detector. Retention times and mass fragmentation signatures of all metabolites were validated using pure standards. To determine the relative metabolite abundance across samples, the peak area for the metabolite of interest was normalized to internal standard and protein content. The mass isotopologue distribution analysis measured the fraction of each metabolite pool that contained every possible number of ^13^C atoms: a metabolite could contain 0, 1, 2, …n ^13^C atoms, where n = the number of carbons in the metabolite. For each metabolite, an informative fragment ion containing all carbons in the parent molecule was analyzed by MATLAB software (MathWorks, CA, USA). The abundance of all mass isotopologues were integrated from m+0 to m+n, where m = the mass of the fragment ion without any ^13^C. The abundance of each mass isotopologue was then corrected mathematically to account for natural abundance isotopes and finally converted into a percentage of the total pool.

### Cell proliferation/growth assay

FASN-WT and FASN-KO H460 cells as well as parental H460 cells were cultured in 6-well plates. Collected cells were stained with 0.4% trypan blue in a 1:1 ratio for 3 minutes. Subsequently, the live (unstained) cells were counted on a hemocytometer. This counting was replicated three times. The proliferation curve was made according to the number of live cells for 7 days. FASN-WT and FASN-KO H460 cells were cultured in attached flat-bottom and non-adherent round-bottom 96-well plates for 48 hours, and the monolayer and spheroid growth were measured by Cell Titer-Glo Assay (Promega, Madison, WI, USA).

Intracellular ATP levels representing cell growth were measured using the Cell Titer-Glo Luminescent Cell Viability Assays (Promega). ATP concentrations were normalized by cell number, and data were expressed as relative X-fold changes versus untreated control cells. H460 FASN-WT and FASN-KO cells (4×10^3^) were seeded in 96-well plates with directly detached treatments of DMSO and 10µM DMNQ for 24 hours to test cell growth. H460 FASN-KO2 cells (4×10^3^) were seeded in 96-well plates with directly detached treatments of DMSO, 10µM DMNQ, 10µM DMNQ plus 200µM CTPi (DMNQ+CTPi), 10µM DMNQ plus 5µM IDH1i (DMNQ+IDH1i), 10µM DMNQ plus 5µM IDH2i, and 10µM DMNQ plus 5µM IDH2i (DMNQ+IDH2i) for 24 hours to test cell growth.

### Immunoblot Analysis

Immunoblot analysis were performed as previously described (Dai et al., 2022). The protein concentrations of cell lysates were determined by the BCA protein assay using bovine serum albumin (BSA) as a standard (Thermo, MA, USA). An equal amount of protein (10μg) per sample was separated on 10% SDS polyacrylamide gels and then transferred onto PVDF membranes (Millipore, MA, USA). The membrane was blocked with 4% non-fat milk in TBST buffer at room temperature for 2 hours, followed by incubation at 4 °C overnight with primary FASN (dilution 1:1000; CST, MA, USA), IDH1 (dilution 1:1000; CST), CTP (dilution 1:500; CST), OXPHOS Cocktail from Abcam (1:1000) and β-actin (dilution 1:4000; CST) antibodies. After washing with TBST three times, the membrane was incubated with secondary goat-anti-rabbit IgG (Bio-Rad Laboratories, CA, USA; dilution 1:3000) or goat anti-mouse IgG (Bio-Rad, dilution 1:3000) conjugated with horseradish peroxidase in 4% non-fat milk for 2 hours at 37°C. Membrane was then washed with TBST three times and incubated with enhanced chemiluminescence Western blotting substrate (Thermo), followed by visualization using an Amersham Imager 680 system (GE Healthcare, MA, USA). The level of β-actin was used as a loading control.

### MitoSOX Red staining assay

For detached mtROS determination, cells were seeded on imaging chambers (Ibidi, WI, USA) with indicated treatments of 24 hours. Then mitochondrial ROS were labeled with MitoSOX Red (5µM; Invitrogen, CA, USA) for 20 min at 37°C. Complete media with different treatments (supplemented with 5% FBS, 4 mM g L-glutamine, and antibiotics) was used for imaging performed on a Zeiss Imager equipped with a N-Achroplan 40 X/0.75 water immersion lens and an AxioCAM MRm digital camera. Images were captured using AxioVision 4.8 and Zeiss Zen software. The images were quantified using Image Pro-premier software 9.3 (Rockville, MD).

### Intracellular NADPH and GSH Measurement

Total cell lysates were used to determine NADPH levels in FASN-WT/-KO cells in 6-cm dish. The quantification of NADPH was assessed in total cell lysates using the NADP/NADPH Assay Kit (Promega) following the manufacturer’s instructions, and intracellular NADPH was normalized to their corresponding protein content. 1×10^4^ of FASN-WT/-KO cells were seeded in 96-well plates overnight. The media was removed, and the total cellular GSH of each cell line was assessed using the GSH-Glo™ Glutathione Assay (Promega) following the manufacturer’s instructions, and intracellular GSH was normalized to their corresponding protein content.

### Seahorse assay

The real-time measurements of OCR of H460-FASN-WT/-KO cells were detected on a Seahorse XF24 Extracellular Flux Analyzer (Agilent Technologies). 1×10^4 cells/well were incubated with XF assay medium (pH 7.4) with 4 mM glutamine and 10 mM glucose, then following compounds were added in sequence: 1.5 µM oligomycin, 2.0 µM carbonyl cyanide 4-(trifluoromethoxy) phenylhydrazone (FCCP), and 1.0 mM rotenone plus antimycin A.

### Metabolic flux analysis (MFA)

Steady-state metabolic fluxes were calculated by combining extracellular flux rates (glucose/ glutamine utilization, lactate/alanine/glutamate secretion) and ^13^C mass isotopologue distributions (MIDs) for citrate, glutamate, fumarate, malate, aspartate, glutamate, and palmitate, using the INCA software package (Young, 2014), which applies an elementary metabolite unit framework to efficiently simulate MIDs (Antoniewicz et al., 2007, Young et al., 2008). We developed reaction networks describing the stoichiometry and carbon transitions of central carbon metabolism, with assumptions as summarized below:

(1) During the experiments, cells are at a metabolic steady state.
(2) ^13^CO_2_ produced during oxidation reactions is not reincorporated via carboxylation reactions.
(3) Cells are given 20 hours to metabolize ^13^C substrates. After 20 hours, it is assumed that the isotopic labeling has reached a steady state.
(4) The metabolites succinate and fumarate are symmetrical and their metabolism through the TCA cycle does not produce a particular orientation.
(5) The metabolites pyruvate, acetyl-CoA, citrate, α-KG, malate, fumarate, and oxaloacetate are metabolically active in both the cytosol and mitochondrion. Malate and α-KG are allowed to freely mix between the compartments.
(6) During the extraction process, intracellular pools of metabolites are homogenized. Therefore, GC-MS analysis of the isotopic enrichment of these metabolites reflects the mixture of distinct metabolic pools. By employing the INCA platform to perform MFA, it is possible to extract meaningful information from these mixed pools. To do this, the model employs parameters to account for the mixing of mitochondrial and cytosolic metabolites.
(7) Parallel labeling data from cultures fed [U-^13^C]glucose and [U-^13^C]glutamine (FASN-WT and FASN-KO2 H460 cells) were used to simultaneously fit the same network model to estimate intracellular fluxes. The labeling data from cultures fed [U-^13^C]glucose and [U-^13^C]glutamine (FASN-WT and FASN-KO2 H460 cells) were used to simultaneously fit the same network model to estimate intracellular fluxes. To ensure that a global minimum of fluxes was identified, flux estimations were initiated from random values and repeated a minimum of 50 times. A chi-square test was applied to test goodness-of-fit, and accurate 95% confidence intervals were calculated by assessing the sensitivity of the sum of squared residuals to flux parameter variations. Table S1 contains the degrees of freedom and sum-of-squared residuals (SSR) for the best fit model and the lower and upper bounds of 95% confidence intervals for all fluxes.

### Quantification and statistical analysis

Statistical analysis was calculated using Graphpad Prism software (GraphPad, CA, USA) and SAS software (version 9.2, SAS Institute, Cary, NC, USA). Unless indicated, all results shown as mean ± SD of cellular triplicates obtained from one representative experiment as specified in each figure legend. To assess the significance of differences among cultures and conditions, a two-tailed Welch’s unequal variances t-test was used to assess the significance between two groups. For three or more groups, a one-way or two-way ANOVA followed by Dunnett’s multiple comparisons test was performed. A *P*-value less than 0.05 was considered significant. All statistical tests were two-tailed.

## Supplemental information

Supplemental information can be found online.

## Acknowledgements

This work was supported by the City of Hope Medical Center Start-up and P30CA033572. R.J.D. is supported by National Cancer Institute Grant R35CA22044901 and CPRIT RP180778. Q.A.W. was supported by National Institutes of Health Grants R01HD096152, R01AG063854, and R01DK128907, as well as the American Diabetes Association Junior Faculty Development Award 1-19-JDF-023. We thank Professor Irfan Lodhi for sharing the pLJM1-Flag-FASN plasmid. Schematics included in the figures were created using BioRender.com.

## Author concentrations

W.T.D. and L.J. conceptualized and designed the study. W.T.D. performed all the experiments and data analysis. L.J. helped with the experiments on fatty acid synthesis and hypoxia culture. Z.C.W. helped with metabolic flux modeling. G.W. helped with flow cytometry. R.J.D. helped revise this manuscript. W.T.D. and L.J. interpreted the data, wrote and revised the manuscript. L.J. and Q.A.W. were responsible for the funding and supervised the study. All authors approved to submit this version for publication.

## Figure legends

**Figure S1.**
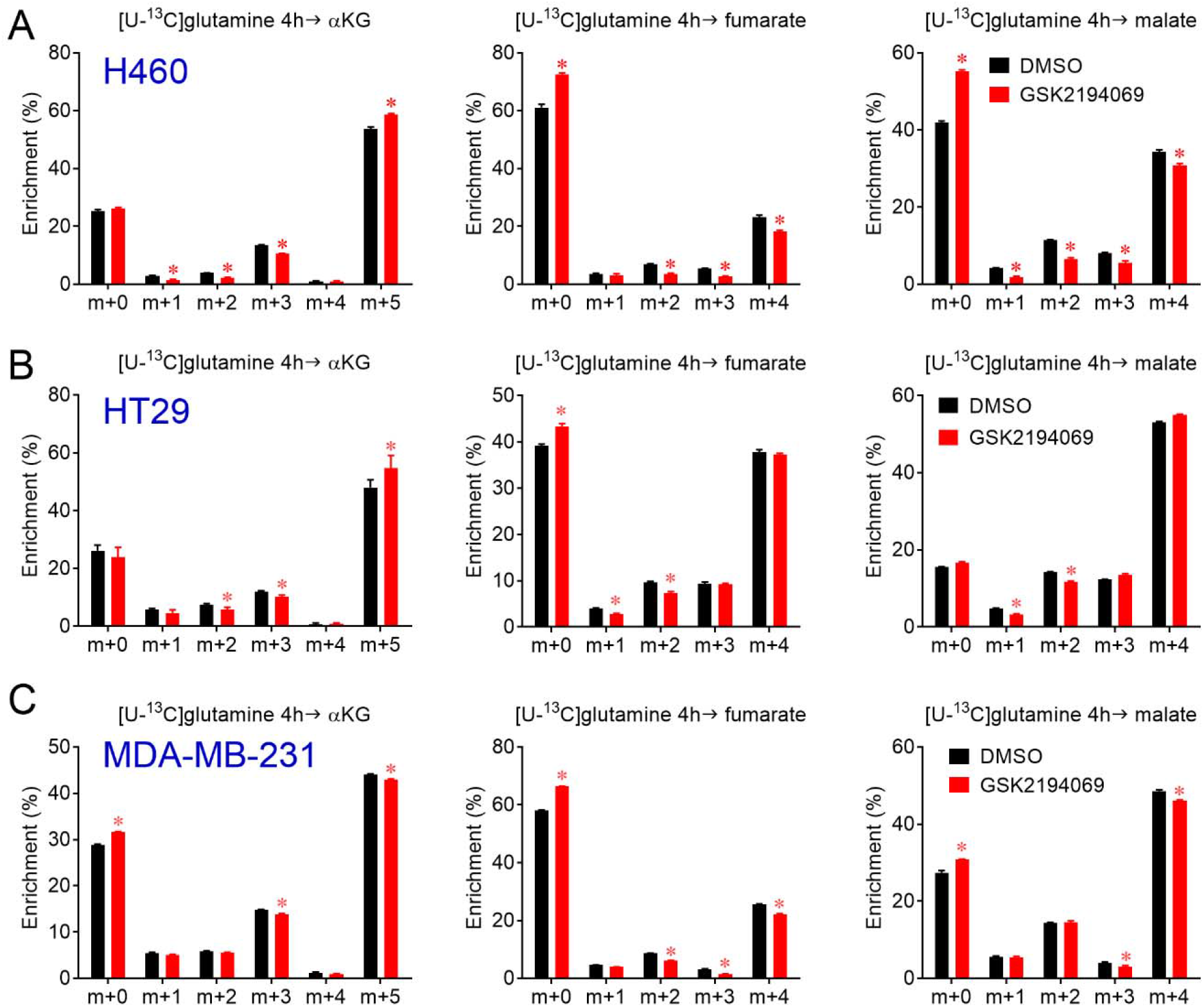
Glutamine metabolism of multiple cancer cells treated with FASN inhibitor (Supporting data for Figure 1). Lung cancer H460 (A), colorectal cancer HT29 (B), and breast cancer MDA-MB-231 (C) cell lines were pretreated with DMSO or 50 µM GSK2194069 for 16 hours, and followed by another 4 hours incubation with tracing medium containing [U-^13^C]glutamine. Mass isotopologue analysis of alpha-ketoglutarate (αKG), fumarate, and malate enrichment were analyzed by GC-MS. The mean ± SD error bars are displayed. *P* values are derived from a two-tailed Welch’s unequal variances t-test between DMSO- and 50 µM GSK2194069-treated groups in each cell line (\**P* < 0.05 compared to DMSO treatment). All experiments were repeated 3 times or more, and n = 3 independent samples.

**Figure S2.**
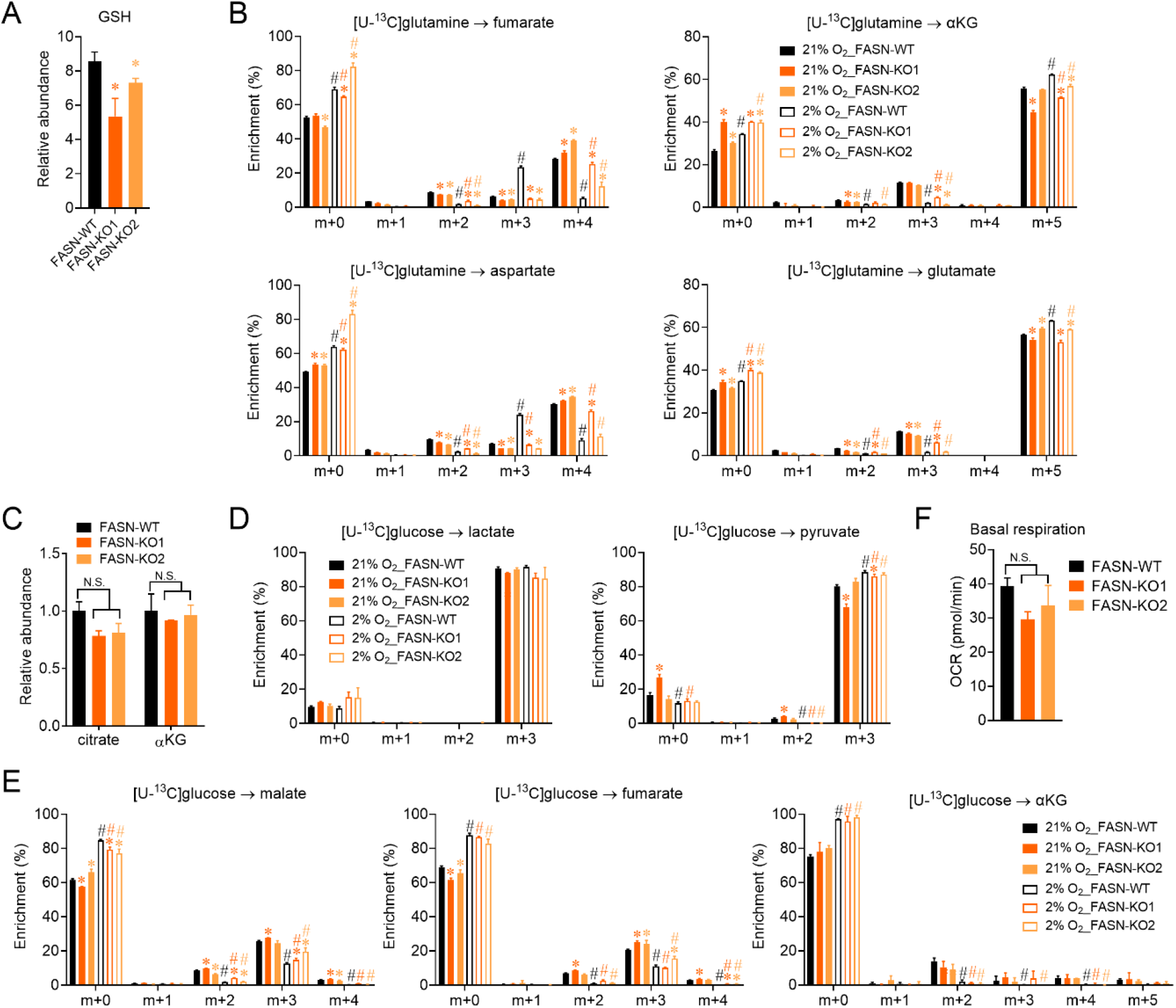
Glutamine and glucose metabolism of FASN-WT and FASN-KO H460 cells under normoxia and hypoxia (Supporting data for Figure 2). (A) The relative abundance of GSH in FASN-WT and FASN-KO H460 cells (n = 3). (B) Mass isotopologue analysis of fumarate, aspartate, αKG, and glutamate in FASN-WT and FASN-KO H460 cells incubated with [U-^13^C]glutamine tracing medium for 4 hours under normoxia and hypoxia (21% *vs*. 2% O_2_). (C) The relative abundance of αKG and citrate in FASN-WT and FASN-KO H460 cells (n = 3). (D) Mass isotopologue analysis of lactate and pyruvate in FASN-WT and FASN-KO H460 cells incubated with [U-^13^C]glucose tracing medium for 4 hours under normoxia and hypoxia. (E) Mass isotopologue analysis of malate, fumarate, and αKG in FASN-WT and FASN-KO H460 cells incubated with [U-^13^C]glucose tracing medium for 4 hours under normoxia and hypoxia. (F) The basal respiration of FASN-WT and FASN-KO H460 cells (n = 6–7). A, C and F: The mean ± SD error bars are displayed. *P* values are derived from a one-way ANOVA followed by Dunnett’s multiple comparisons test for the three cell lines. *P* values compared to the FASN-WT group: N.S. represents no significance (*P* > 0.05). (\**P* < 0.05 compared to the FASN-WT group). B, D, and E: The mean ± SD error bars are displayed. *\*P* values are derived from a one-way ANOVA followed by Dunnett’s multiple comparisons test for the three cell lines under normoxia or hypoxia (*\*P* < 0.05 compared to 21% O2_FASN-WT group or 2% O2_FASN-WT group). *^#^P* values are derived from a two-tailed Welch’s unequal variances t-test between two groups in each cell line (*^#^P* < 0.05 comparing the 21% O2 group to the 2% O2 group). All experiments were repeated 3 times or more, and n = 3 independent samples.

**Figure S3.**
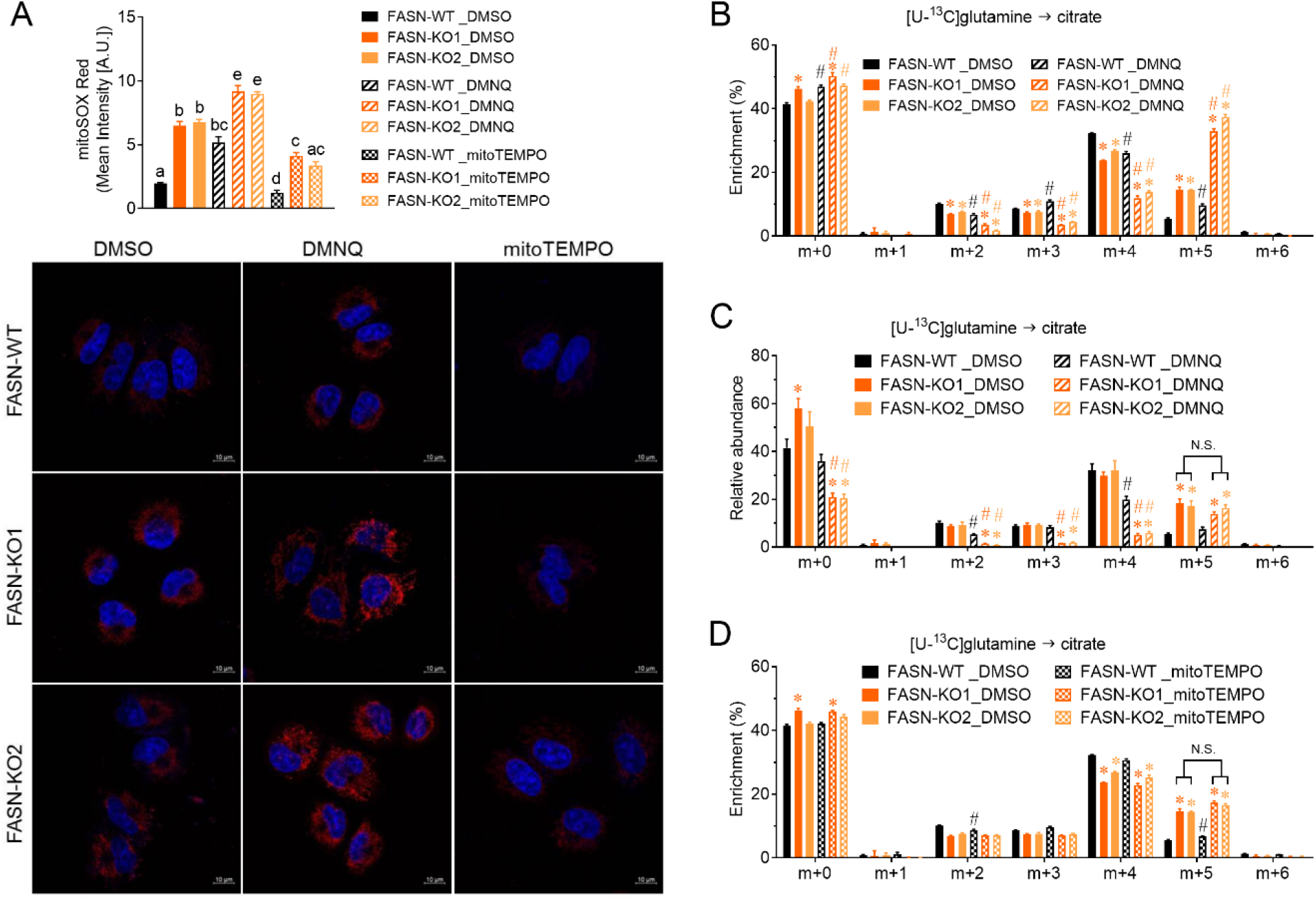
Mitochondrial ROS and reductive carboxylation co-exist in FASN-KO cells (Supporting data for Figure 2). (A) Mitochondrial ROS levels measured by mitoSOX Red staining in FASN-WT and FASN-KO cells treated with DMSO, 10 μM DMNQ, and 100 μM mitoTEMPO (n = 4). The scale bar represents 10 μm. The mean ± SD error bars are displayed. *P* values are derived from two-way ANOVA followed by Dunnett’s multiple comparisons test for the 9 groups with different treatments in all three cell lines. *P* values with different lowercase letters among groups represent significance (< 0.05), and *P* values containing the same lowercase letter among groups represent no significance (> 0.05). (B) Mass isotopologue analysis of citrate enrichment in FASN-WT and FASN-KO cells incubated with [U-^13^C]glutamine tracing medium containing 10 μM DMNQ for 4 hours. (C) Relative abundance of the citrate isotopologues in panel B. (D) Mass isotopologue analysis of citrate enrichment in FASN-WT and FASN-KO cells incubated with [U-^13^C]glutamine tracing medium containing 100 μM mitoTEMPO for 4 hours. B-D: The mean ± SD error bars are displayed. *\*P* values are derived from a one-way ANOVA followed by Dunnett’s multiple comparisons test for the three cell lines with DMSO/DMNQ/ mitoTEMPO treatment (*\*P* < 0.05 compared to FASN-WT_DMSO or FASN-WT_DMNQ or FASN-WT_mitoTEMPO). *^#^P* values are derived from a two-tailed Welch’s unequal variances t-test between two groups in each cell line under different treatment conditions (*^#^P* < 0.05 compared DMNQ/mitoTEMPO groups to DMSO groups). N.S. represents no significance (*P* > 0.05). All experiments were repeated 3 times or more, and n = 3 independent samples.

**Figure S4.**
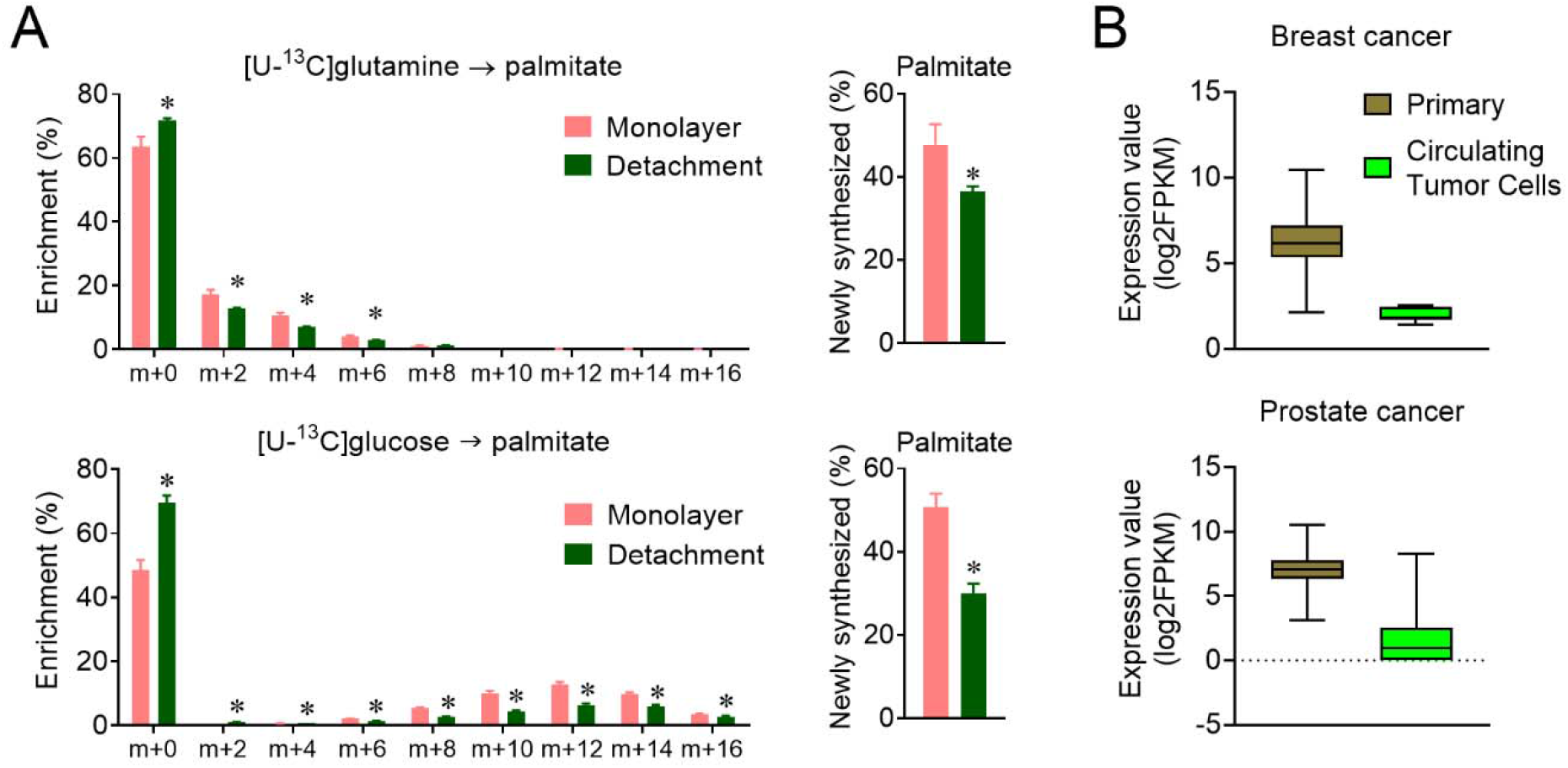
Palmitate synthesis in monolayer and spheroid (Supporting data for Figure 4). (A) The mass isotopologue analysis of palmitate enrichment in H460 monolayer and spheroid culture incubated with [U-^13^C]glutamine tracing medium and [U-^13^C]glucose tracing medium for 20 hours. The Palmitate synthesis rate is calculated from the palmitate enrichment data. *P* values are derived from a two-tailed Welch’s unequal variances t-test between monolayer and spheroid-cultured H460 cells (\**P* < 0.05 compared to monolayer culture). These are re-analyzed data in the previous H460 spheroid study (Jiang et al., 2016). (B) Boxplot of FASN gene expression levels in primary cancer cells and circulating tumor cells (CTC) from patients with breast cancer and prostate cancer. The boxplot was made from the data downloaded from the gene expression database of CTC (ctcRbase) (Miyamoto et al., 2015, Yu et al., 2014).

**Figure S5.**
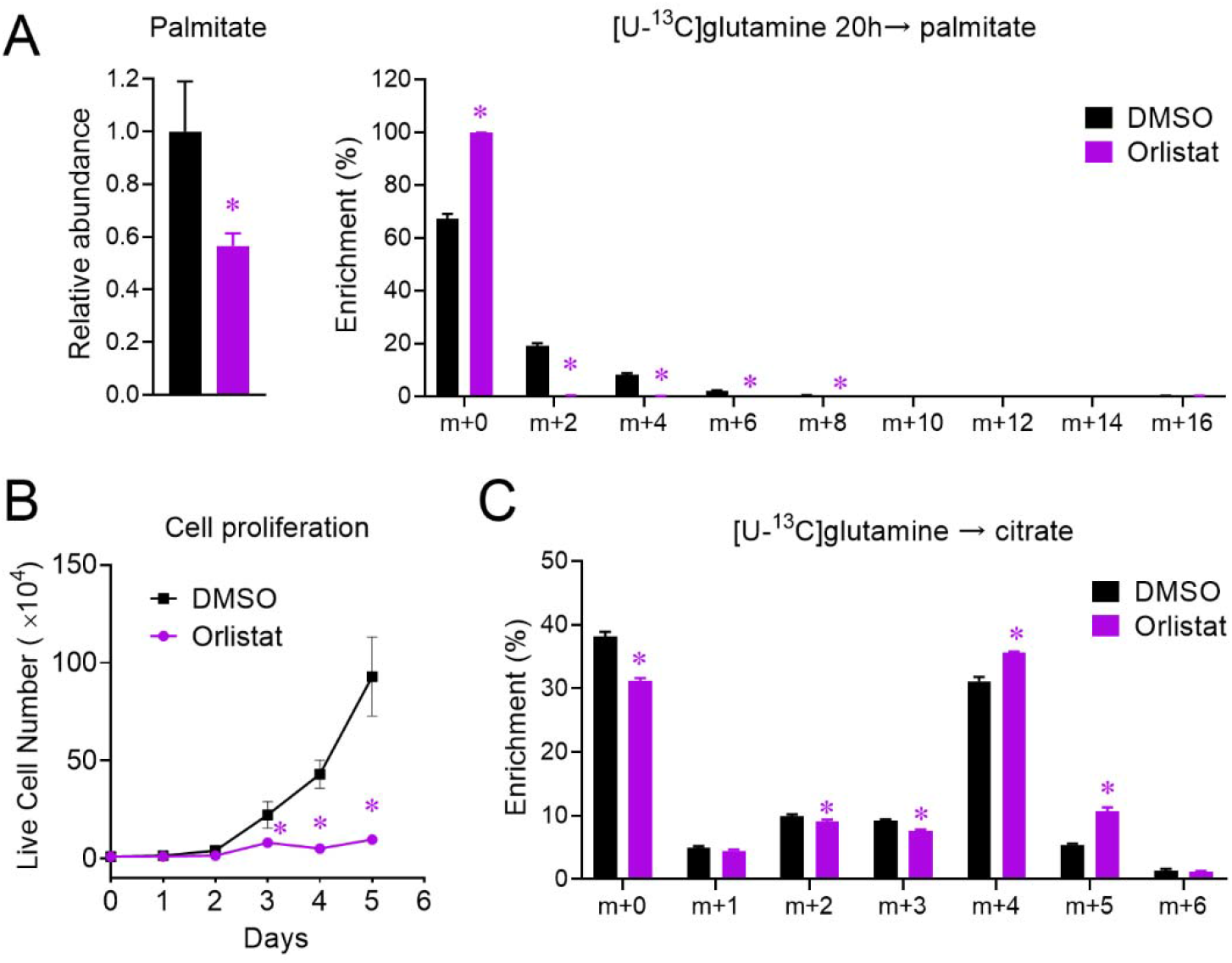
Orlistat induces reductive carboxylation in H460 cells. (A) The relative abundance and mass isotopologue analysis of palmitate in H460 cells incubated with [U-^13^C]glutamine tracing medium containing 50 μM Orlistat for 20 hours. (B) Cell proliferation of H460 cells incubated with DMSO and 50 μM Orlistat. (C) Mass isotopologue analysis of citrate enrichment in H460 cells pretreated with 50 μM Orlistat for 16 hours and incubated with [U-^13^C]glutamine tracing medium containing the corresponding treatments for another 4 hours. A-C: The mean ± SD error bars are displayed. *P* values are derived from a two-tailed Welch’s unequal variances t-test between DMSO- and Orlistat-treated groups (\**P* < 0.05 compared to DMSO treatment). All experiments were repeated 3 times or more, and n = 3 independent samples.

**Table S1.**
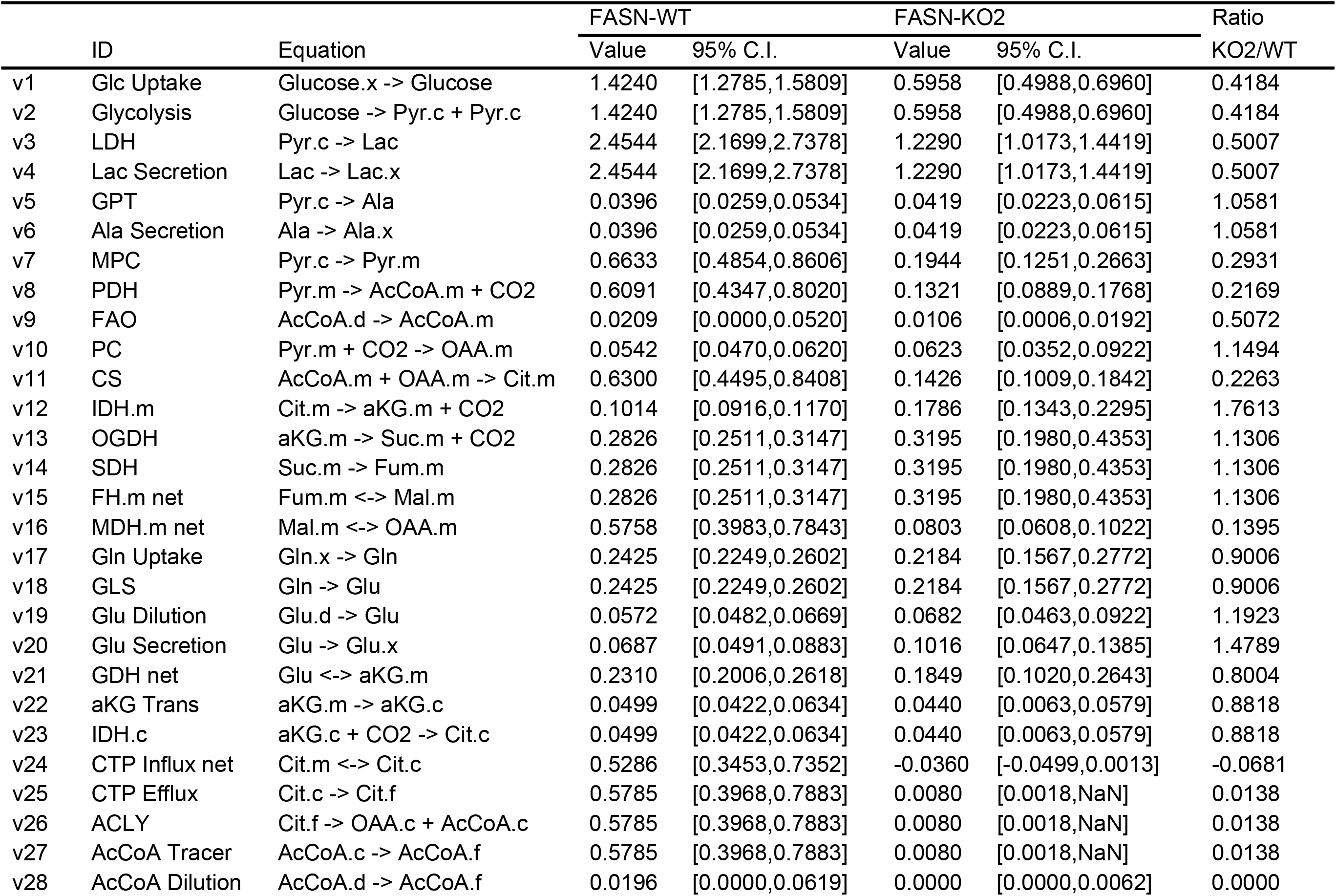
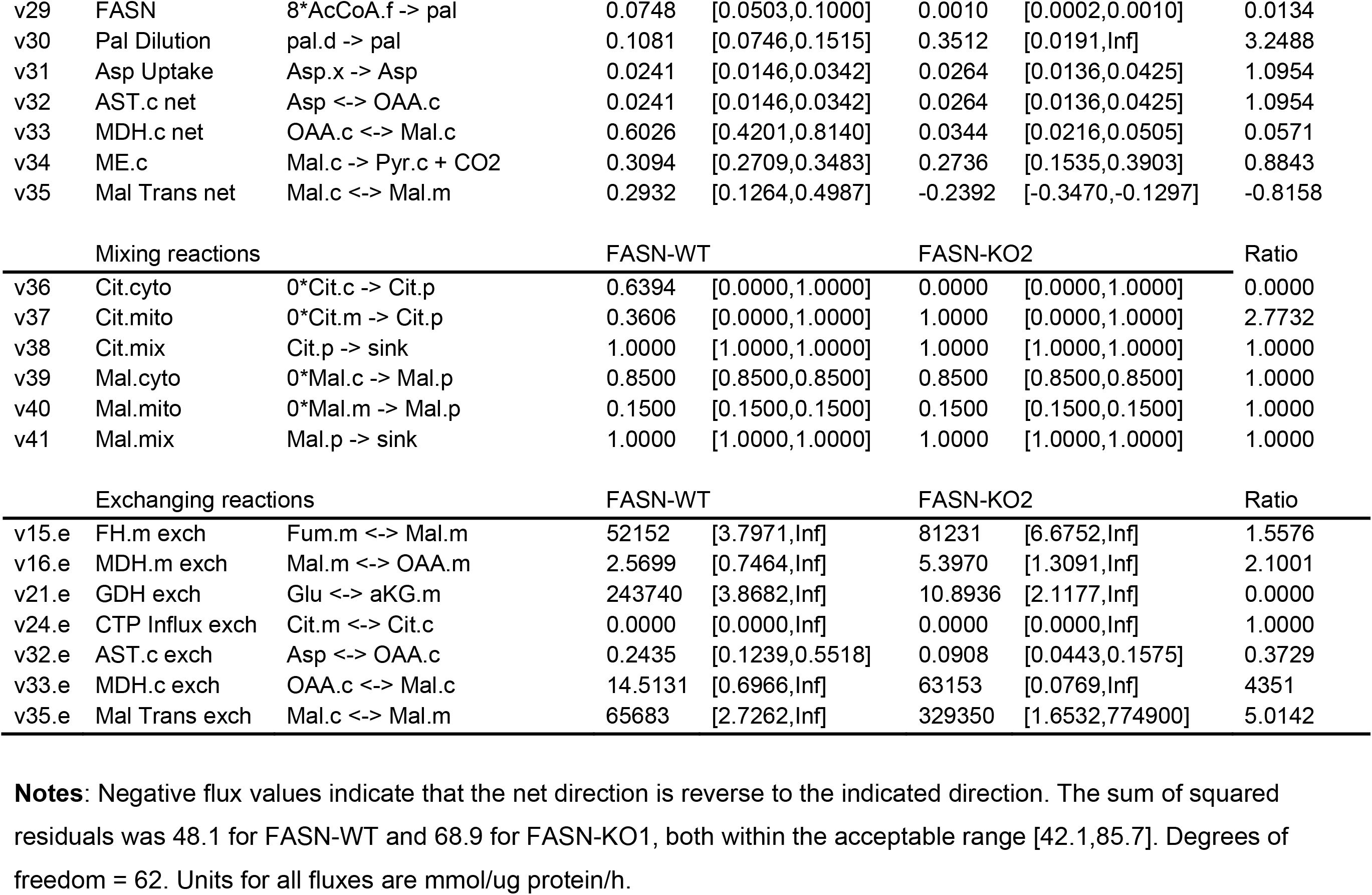
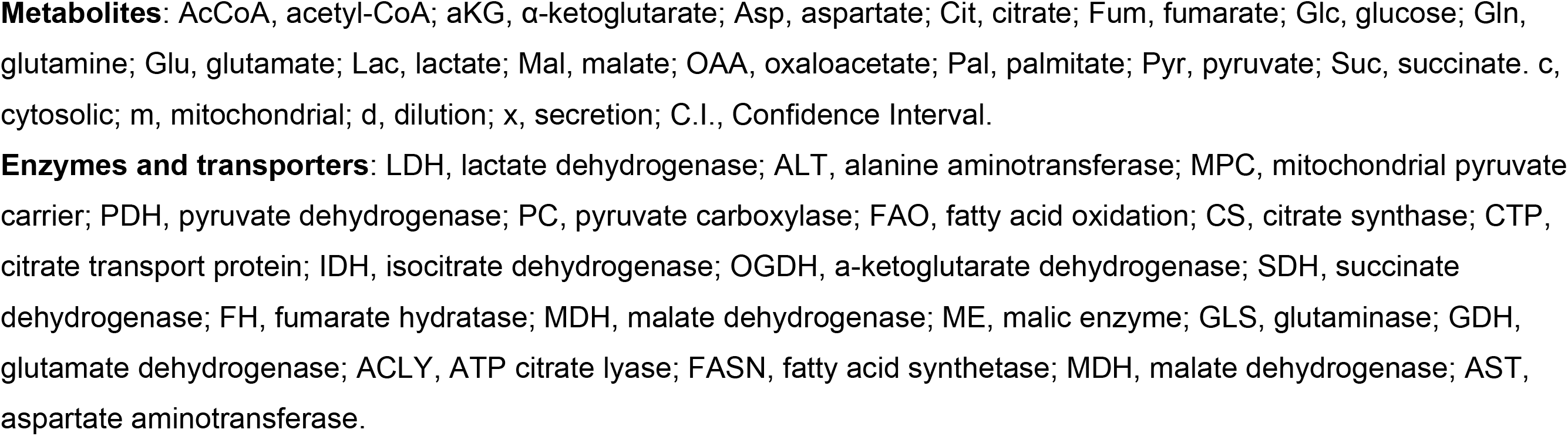
Simulated metabolic fluxes in FASN-WT and FASN-KO H460 cells.

## References

Alli PM, Pinn ML, Jaffee EM, McFadden JM, Kuhajda FP (2005) Fatty acid synthase inhibitors are chemopreventive for mammary cancer in neu-N transgenic mice. Oncogene 24: 39–46

Altman BJ, Stine ZE, Dang CV (2016) From Krebs to clinic: glutamine metabolism to cancer therapy. Nature reviews Cancer 16: 619–634

Antoniewicz MR, Kelleher JK, Stephanopoulos G (2007) Elementary metabolite units (EMU): a novel framework for modeling isotopic distributions. Metabolic engineering 9: 68–86

Arnold PK, Jackson BT, Paras KI, Brunner JS, Hart ML, Newsom OJ, Alibeckoff SP, Endress J, Drill E, Sullivan LB (2022) A non-canonical tricarboxylic acid cycle underlies cellular identity. Nature 603: 477–481

Bandyopadhyay S, Zhan R, Wang Y, Pai SK, Hirota S, Hosobe S, Takano Y, Saito K, Furuta E, Iiizumi M, Mohinta S, Watabe M, Chalfant C, Watabe K (2006) Mechanism of apoptosis induced by the inhibition of fatty acid synthase in breast cancer cells. Cancer research 66: 5934–40

Batchuluun B, Pinkosky SL, Steinberg GR (2022) Lipogenesis inhibitors: therapeutic opportunities and challenges. Nature Reviews Drug Discovery 21: 283–305

Bergers G, Fendt S-M (2021) The metabolism of cancer cells during metastasis. Nature Reviews Cancer 21: 162–180

Bueno MJ, Jimenez-Renard V, Samino S, Capellades J, Junza A, López-Rodríguez ML, Garcia-Carceles J, Lopez-Fabuel I, Bolaños JP, Chandel NS, Yanes O, Colomer R, Quintela-Fandino M (2019) Essentiality of fatty acid synthase in the 2D to anchorage-independent growth transition in transforming cells. Nature communications 10: 5011

Dai WT, Wang G, Chwa J, Oh ME, Abeywardana T, Yang YZ, Wang QA, Jiang L (2020) Mitochondrial division inhibitor (mdivi-1) decreases oxidative metabolism in cancer. British journal of cancer 122: 1288–1297

Dai WT, Wang ZC, Wang QA, Chan D, Jiang L (2022) Metabolic reprogramming in the OPA1-deficient cells. Cellular and molecular life sciences : CMLS 79: 517

Droin C, Kholtei JE, Bahar Halpern K, Hurni C, Rozenberg M, Muvkadi S, Itzkovitz S, Naef F (2021) Space-time logic of liver gene expression at sub-lobular scale. Nature metabolism 3: 43–58

Faubert B, Solmonson A, DeBerardinis RJ (2020) Metabolic reprogramming and cancer progression. Science 368: eaaw5473

Fendt SM, Bell EL, Keibler MA, Olenchock BA, Mayers JR, Wasylenko TM, Vokes NI, Guarente L, Vander Heiden MG, Stephanopoulos G (2013) Reductive glutamine metabolism is a function of the α-ketoglutarate to citrate ratio in cells. Nature communications 4: 2236

Ferraro GB, Ali A, Luengo A, Kodack DP, Deik A, Abbott KL, Bezwada D, Blanc L, Prideaux B, Jin X, Possada JM, Chen J, Chin CR, Amoozgar Z, Ferreira R, Chen I, Naxerova K, Ng C, Westermark AM, Duquette M et al. (2021) Fatty acid synthesis is required for breast cancer brain metastasis. Nature cancer 2: 414–428

Fischer GM, Jalali A, Kircher DA, Lee W-C, McQuade JL, Haydu LE, Joon AY, Reuben A, de Macedo MP, Carapeto FC (2019) Molecular profiling reveals unique immune and metabolic features of melanoma brain metastases. Cancer discovery 9: 628–645

Grassian A, Coloff J, Brugge J (2011) Extracellular matrix regulation of metabolism and implications for tumorigenesis. In Cold Spring Harbor symposia on quantitative biology, pp 313-324. Cold Spring Harbor Laboratory Press

Hanahan D (2022) Hallmarks of Cancer: New Dimensions. Cancer Discovery 12: 31&<otherinfo>-46&</otherinfo>

Hardwicke MA, Rendina AR, Williams SP, Moore ML, Wang L, Krueger JA, Plant RN, Totoritis RD, Zhang G, Briand J, Burkhart WA, Brown KK, Parrish CA (2014) A human fatty acid synthase inhibitor binds β-ketoacyl reductase in the keto-substrate site. Nature chemical biology 10: 774–9

Hlouschek J, Hansel C, Jendrossek V, Matschke J (2018) The mitochondrial citrate carrier (SLC25A1) sustains redox homeostasis and mitochondrial metabolism supporting radioresistance of cancer cells with tolerance to cycling severe hypoxia. Frontiers in oncology 8: 170

Jiang L, Deberardinis R, Boothman DA (2015a) The cancer cell ‘energy grid’: TGF-β1 signaling coordinates metabolism for migration. Molecular & cellular oncology 2: e981994

Jiang L, Shestov AA, Swain P, Yang C, Parker SJ, Wang QA, Terada LS, Adams ND, McCabe MT, Pietrak B, Schmidt S, Metallo CM, Dranka BP, Schwartz B, DeBerardinis RJ (2016) Reductive carboxylation supports redox homeostasis during anchorage-independent growth. Nature 532: 255–8

Jiang L, Wang H, Li J, Fang X, Pan H, Yuan X, Zhang P (2014) Up-regulated FASN expression promotes transcoelomic metastasis of ovarian cancer cell through epithelial-mesenchymal transition. International journal of molecular sciences 15: 11539–54

Jiang L, Xiao L, Sugiura H, Huang X, Ali A, Kuro-o M, Deberardinis RJ, Boothman DA (2015b) Metabolic reprogramming during TGFβ1-induced epithelial-to-mesenchymal transition. Oncogene 34: 3908–16

Knowles LM, Axelrod F, Browne CD, Smith JW (2004) A fatty acid synthase blockade induces tumor cell-cycle arrest by down-regulating Skp2. The Journal of biological chemistry 279: 30540–5

Kridel SJ, Axelrod F, Rozenkrantz N, Smith JW (2004) Orlistat is a novel inhibitor of fatty acid synthase with antitumor activity. Cancer research 64: 2070&<otherinfo>-5&</otherinfo>

Li Z, Ji BW, Dixit PD, Tchourine K, Lien EC, Hosios AM, Abbott KL, Rutter JC, Westermark AM, Gorodetsky EF, Sullivan LB, Vander Heiden MG, Vitkup D (2022) Cancer cells depend on environmental lipids for proliferation when electron acceptors are limited. Nature metabolism 4: 711–723

Menendez JA, Lupu R (2007) Fatty acid synthase and the lipogenic phenotype in cancer pathogenesis. Nature reviews Cancer 7: 763–77

Metallo CM, Gameiro PA, Bell EL, Mattaini KR, Yang J, Hiller K, Jewell CM, Johnson ZR, Irvine DJ, Guarente L, Kelleher JK, Vander Heiden MG, Iliopoulos O, Stephanopoulos G (2011) Reductive glutamine metabolism by IDH1 mediates lipogenesis under hypoxia. Nature 481: 380–4

Miyamoto DT, Zheng Y, Wittner BS, Lee RJ, Zhu H, Broderick KT, Desai R, Fox DB, Brannigan BW, Trautwein J (2015) RNA-Seq of single prostate CTCs implicates noncanonical Wnt signaling in antiandrogen resistance. Science 349: 1351–1356

Mullen AR, Wheaton WW, Jin ES, Chen PH, Sullivan LB, Cheng T, Yang Y, Linehan WM, Chandel NS, DeBerardinis RJ (2011) Reductive carboxylation supports growth in tumour cells with defective mitochondria. Nature 481: 385&<otherinfo>-8&</otherinfo>

Palmieri F, Scarcia P, Monné M (2020) Diseases caused by mutations in mitochondrial carrier genes SLC25: a review. Biomolecules 10: 655

Pavlova NN, Zhu J, Thompson CB (2022) The hallmarks of cancer metabolism: Still emerging. Cell metabolism

Röhrig F, Schulze A (2016) The multifaceted roles of fatty acid synthesis in cancer. Nature reviews Cancer 16: 732–749

Ran FA, Hsu PD, Wright J, Agarwala V, Scott DA, Zhang F (2013) Genome engineering using the CRISPR-Cas9 system. Nat Protoc 8: 2281–2308

Schafer ZT, Grassian AR, Song L, Jiang Z, Gerhart-Hines Z, Irie HY, Gao S, Puigserver P, Brugge JS (2009) Antioxidant and oncogene rescue of metabolic defects caused by loss of matrix attachment. Nature 461: 109–13

Seguin F, Carvalho M, Bastos D, Agostini M, Zecchin K, Alvarez-Flores MP, Chudzinski-Tavassi AM, Coletta R, Graner E (2012) The fatty acid synthase inhibitor orlistat reduces experimental metastases and angiogenesis in B16-F10 melanomas. British journal of cancer 107: 977&<otherinfo>-9&</otherinfo>87

Sies H, Belousov VV, Chandel NS, Davies MJ, Jones DP, Mann GE, Murphy MP, Yamamoto M, Winterbourn C (2022) Defining roles of specific reactive oxygen species (ROS) in cell biology and physiology. Nature reviews Molecular cell biology

Spinelli JB, Haigis MC (2018) The multifaceted contributions of mitochondria to cellular metabolism. Nature cell biology 20: 745–754

Tan M, Mosaoa R, Graham GT, Kasprzyk-Pawelec A, Gadre S, Parasido E, Catalina-Rodriguez O, Foley P, Giaccone G, Cheema A, Kallakury B, Albanese C, Yi C, Avantaggiati ML (2020) Inhibition of the mitochondrial citrate carrier, Slc25a1, reverts steatosis, glucose intolerance, and inflammation in preclinical models of NAFLD/NASH. Cell Death & Differentiation 27: 2143–2157

Tasdogan A, Ubellacker JM, Morrison SJ (2021) Redox regulation in cancer cells during metastasis. Cancer Discovery 11: 2682–2692

Valastyan S, Weinberg Robert A (2011) Tumor Metastasis: Molecular Insights and Evolving Paradigms. Cell 147: 275–292

Wang Z, Ning T, Song A, Rutter J, Wang QA, Jiang L (2020) Chronic cold exposure enhances glucose oxidation in brown adipose tissue. EMBO reports 21: e50085

Wise DR, Ward PS, Shay JE, Cross JR, Gruber JJ, Sachdeva UM, Platt JM, DeMatteo RG, Simon MC, Thompson CB (2011) Hypoxia promotes isocitrate dehydrogenase-dependent carboxylation of α-ketoglutarate to citrate to support cell growth and viability. Proceedings of the National Academy of Sciences of the United States of America 108: 19611–6

Yang C, Ko B, Hensley CT, Jiang L, Wasti AT, Kim J, Sudderth J, Calvaruso MA, Lumata L, Mitsche M, Rutter J, Merritt ME, DeBerardinis RJ (2014) Glutamine oxidation maintains the TCA cycle and cell survival during impaired mitochondrial pyruvate transport. Molecular cell 56: 414-24

Yoo H, Antoniewicz MR, Stephanopoulos G, Kelleher JK (2008) Quantifying reductive carboxylation flux of glutamine to lipid in a brown adipocyte cell line. The Journal of biological chemistry 283: 20621–7

Young JD (2014) INCA: a computational platform for isotopically non-stationary metabolic flux analysis. Bioinformatics (Oxford, England) 30: 1333–5

Young JD, Walther JL, Antoniewicz MR, Yoo H, Stephanopoulos G (2008) An elementary metabolite unit (EMU) based method of isotopically nonstationary flux analysis. Biotechnology and bioengineering 99: 686–99

Yu M, Bardia A, Aceto N, Bersani F, Madden MW, Donaldson MC, Desai R, Zhu H, Comaills V, Zheng Z (2014) Ex vivo culture of circulating breast tumor cells for individualized testing of drug susceptibility. science 345: 216–220

Zaytseva YY, Rychahou PG, Gulhati P, Elliott VA, Mustain WC, O’Connor K, Morris AJ, Sunkara M, Weiss HL, Lee EY (2012) Inhibition of fatty acid synthase attenuates CD44-associated signaling and reduces metastasis in colorectal cancer. Cancer research 72: 1504–1517

Zhi J, Mulligan TE, Hauptman JB (1999) Long-term systemic exposure of orlistat, a lipase inhibitor, and its metabolites in obese patients. Journal of clinical pharmacology 39: 41&<otherinfo>-6&</otherinfo>

